# Serum Proteome Profiling Identifies N-Cadherin and C-Met as Early Marker Candidates of Therapeutic Response to Neoadjuvant Chemotherapy in Breast Cancer

**DOI:** 10.1101/2024.05.24.595719

**Authors:** Ines Derya Steenbuck, Miguel Cosenza-Contreras, Klemens Fröhlich, Bettina Mayer, Konrad Kurowski, Tilman Werner, Meike Reinold, Matthias Fahrner, Frank Hause, Adrianna Seredynska, Tobias Feilen, Andrea Ritter, Armelle Guénégou-Arnoux, Martin L. Biniossek, Daniela Weiss, Claudia Nöthling, Markus Jäger, Thalia Erbes, Oliver Schilling

## Abstract

Breast cancer remains the most common cancer in women worldwide. Neoadjuvant chemotherapy (NACT) is often preferred to adjuvant chemotherapy to achieve tumour shrinkage, monitor response to therapy and facilitate surgical removal in the absence of metastases. In addition, there is strong evidence that pathological complete remission (pCR) is associated with prolonged survival. In this study, we sought to identify candidate markers that signal response or resistance to therapy. We present a retrospective longitudinal serum proteomic study of 22 breast cancer patients (11 with pCR and 11 with non-pCR) matched with 21 healthy controls. Serum was analysed by LC-MS/MS after depletion of abundant proteins by immunoaffinity, trypsinisation, isobaric labelling and fractionation by reversed-phase HPLC. We observed an inverse behaviour of the serum proteins c-Met and N-cadherin after the second cycle of chemotherapy with a high predictive value (AUC 0.93). More pronounced changes were observed after the 6th cycle of NACT, with significant changes in the intensity of the proteins contactin-1, centrosomal protein, sex hormone-binding globuline and cholinesterase. Our study highlights the possibility of monitoring response to NACT using serum as a liquid biopsy.

## 1. Introduction

One in eight women in Germany will be diagnosed with breast cancer during her lifetime (incidence of 112.6 women per 100,000).(1,2). Although mortality rates have fallen in recent years, there is still a standardised death rate of 22.8. After cardiovascular diseases, breast cancer is the fifth most common cause of death in Germany. Early detection is the most important factor in reducing mortality, as advanced stages of cancer greatly reduce the 5-year survival rate (3–5). Up to now, five molecular subtypes of invasive breast cancer are being distinguished in the clinical routine, known as Luminal A-like, Luminal B-like (Her2/neu-negative), Luminal B-like (Her2/neu-positive), Her2/neu-positive (hormone receptor negative) and triple-negative breast cancer (TNBC).

In the case of locoregional primary disease, surgery and chemotherapy are required to minimise the possibility of recurrence. Nowadays, systemic cytostatic therapy is recommended for all breast cancer subtypes except luminal A-like. In terms of overall survival, NACT is equivalent to adjuvant chemotherapy. (6,7). NACT can lead to an improvement of operability, increased rates of breast-conserving surgery, and in case of patients with TNBC and pathological complete remission (pCR), to improvement of long-term clinical prognosis (8,9). A response to NACT is therefore essential to achieve a pathological complete remission (10). The standard chemotherapy for non-metastatic breast cancer is a combination of four cycles of an anthracycline with cyclophosphamide (AC), followed by 12 doses, four cycles of a taxol derivative as paclitaxel (P). In recent years, the interval between AC cycles has typically been three weeks (q3w). A dose-dense interval of two weeks (q2w) was only used in high-risk patients. However, the use of dose-dense AC cycles has shown an improved response compared to three-week cycle intervals, so the current recommendation is a dose-dense AC schedule regardless of nodal status (7,11–13). Furthermore, the addition of carboplatin (Cp) to the treatment regimen of TNBC patients has been shown to be beneficial, and the standard of care for TNBC currently includes four cycles of AC followed by 12 doses of P combined with Cp every 3 weeks (14). In this study, patients received only AC dose-dense or conventional regimens with epirubicine and cyclophosphamide followed by four cycles of paclitaxel. Carboplatin was not used.

NACT offers the opportunity to monitor response as an in vivo sensitivity test to adjust the treatment regimen. NACT response is monitored using imaging techniques such as ultrasound or MRI. Early response (reduction in tumour size) to chemotherapy has been shown to be a favourable marker for achieving pCR (15). However, tumour shrinkage is not a reliable indicator of pCR. Therefore, additional prognostic markers for pCR are of central importance. Because of their low invasiveness, associated low risk of complications and low resource commitment, liquid biopsies offer an elegant diagnostic method for therapy validation at short intervals. Serum is a well-known medium for targeted liquid biopsies. It is also a rich source for the study of non-invasive biomarkers, including proteins, from just a few microlitres of serum. Due to its heterogeneous composition, the exploratory approach in proteomics using liquid chromatography-tandem mass spectrometry (LC-MS/MS) remains a challenge. In the last decade, parameters such as depletion of high-abundance proteins, fractionation or labelling have been adapted to improve the depth of detection in bottom-up proteomics. As a result, the maximum number of proteins identified is also very broad, ranging from 69 (method: immunodepletion by Seppro® IgY14 column, followed by tryptic digestion, filtration, analysis by nano ultraperformance liquid chromatography coupled to MS) to 4,631 (method: immunodepletion by Seppro® IgY14/Supermix columns, followed by LysC-Trypsin Co digestion, iTRAQ labelling, fractionation, LC-MS/MS) (16–18).

To our knowledge, this is the first study to investigate the serum protein imprint of NACT in BC in a longitudinal, patient-matched manner. We compared the patient-matched serum proteome over time, distinguishing between NACT responders, defined as patients with pCR (CR), and non-responders, defined as patients with non-pCR (NR). Here we describe several proteins that show an inverse behaviour compared to CR versus NR at different time points during NACT. Already after two cycles of chemotherapy, we identified two serum proteins (N-cadherin, c-MET) with a high predictive value (AUC = 0.93) based on their time-dependent inter-individual change. In addition, we compared the serum proteomic fingerprint of BC patients before NACT with a control group (Ctrl). We report an activated immune response by upregulation of proteins involved in the complement system and downregulation of immunoglobulin fragments in BC patients. Our findings may lead to potential prognostic serum biomarkers for BC patients treated with NACT.

## 2. Materials and Methods

### 2.1 Ethics statement

The study was approved by the Ethics Committee of the University Medical Center Freiburg (607/16). All patients gave informed consent.

### 2.2 Patient cohort

22 patients were included, of whom 6 were diagnosed with TNBC and 16 with luminal B-like (HER2/neu) breast cancer. Of these patients, 11 had pCR and 11 had non-pCR on histopathological examination after surgery. Serum was collected at three different time points for each patient. In addition, 21 healthy female volunteers were included as a control group. Clinicopathological and NACT details are shown in Table I.

**Table I.** Patients included by response to therapy, subtype, histopathological result, differentiation, age.

| Characteristics |  | pCR patients |  | Non-pCR patients |  | Control group |
| --- | --- | --- | --- | --- | --- | --- |
|  |  | LumB (n=8) | TNBC (n=3) | LumB (n=8) | TNBC (n=3) | (n=21) |
| Age, m+/-SD |  | 54.25 +/-11.85 | 51.33 +/-13.32 | 45.38 +/-11.12 | 51.66 +/-14.29 | 55.48 +/-15.77 |
| Sex | Male | N = 0 | N = 0 | N = 0 | N = 0 | N = 0 |
|  | Female | N = 8 | N = 3 | N = 8 | N = 3 | N = 21 |
| Grade | G3 | 6 | 3 | 3 | 2 | NA |
|  | G2 | 2 | 0 | 5 | 1 |  |
|  | G1 | 0 | 0 | 0 | 0 |  |
| TNM | ypT0 | 8 | 3 | 0 | 0 |  |
|  | ypT1b | 0 | 0 | 1 | 2 |  |
|  | ypT1c | 0 | 0 | 3 | 0 |  |
|  | ypT2 | 0 | 0 | 1 | 1 |  |
|  | ypT2a | 0 | 0 | 1 | 0 |  |
|  | ypT3 | 0 | 0 | 1 | 0 |  |
|  | ypT4b | 0 | 0 | 1 | 0 |  |
| Number of samples per time point | t0 | 8 | 3 | 8 | 3 |  |
|  | t1 | 8 | 3 | 8 | 3 |  |
|  | t2 | 7 | 2 | 8 | 2 |  |

### 2.3 Sample Preparation

In the case of patient sera, the serum gel monovettes were filled by medical staff during routine blood collection just prior to chemotherapy. Blood was collected either through a port system or a venous cannula. Hygiene standards were maintained by disinfection and trained staff. The monovette was regularly filled to maximum capacity and stored at room temperature until further processing. After coagulation, the monovette was centrifuged at 2000g for 10 minutes at 4°C. The cell-free supernatant, the serum, was then transferred to a reaction tube with a safety margin of 2-3 mm and stored at ™20 °C.

Samples were generally randomised and pseudonymised. All reagents were freshly prepared. The method of choice was serum depletion by immunoaffinity (Seppro® IgY14, Sigma-Aldrich) of 14 highly abundant proteins (albumin, IgG, IgA, transferrin, haptoglobin, antitrypsin, fibrinogen, alpha 2-macroglobulin, alpha 1, IgM, apolipoproteins AI and AII, C3, transthyretin) followed by Lys-C and trypsin co-digestion, clean-up by PreOmics®, tandem mass tag (TMT) -labelling, fractionation by reversed-phase HPLC and LC-MS/MS measurement (Figure 1).

**Figure 1.**
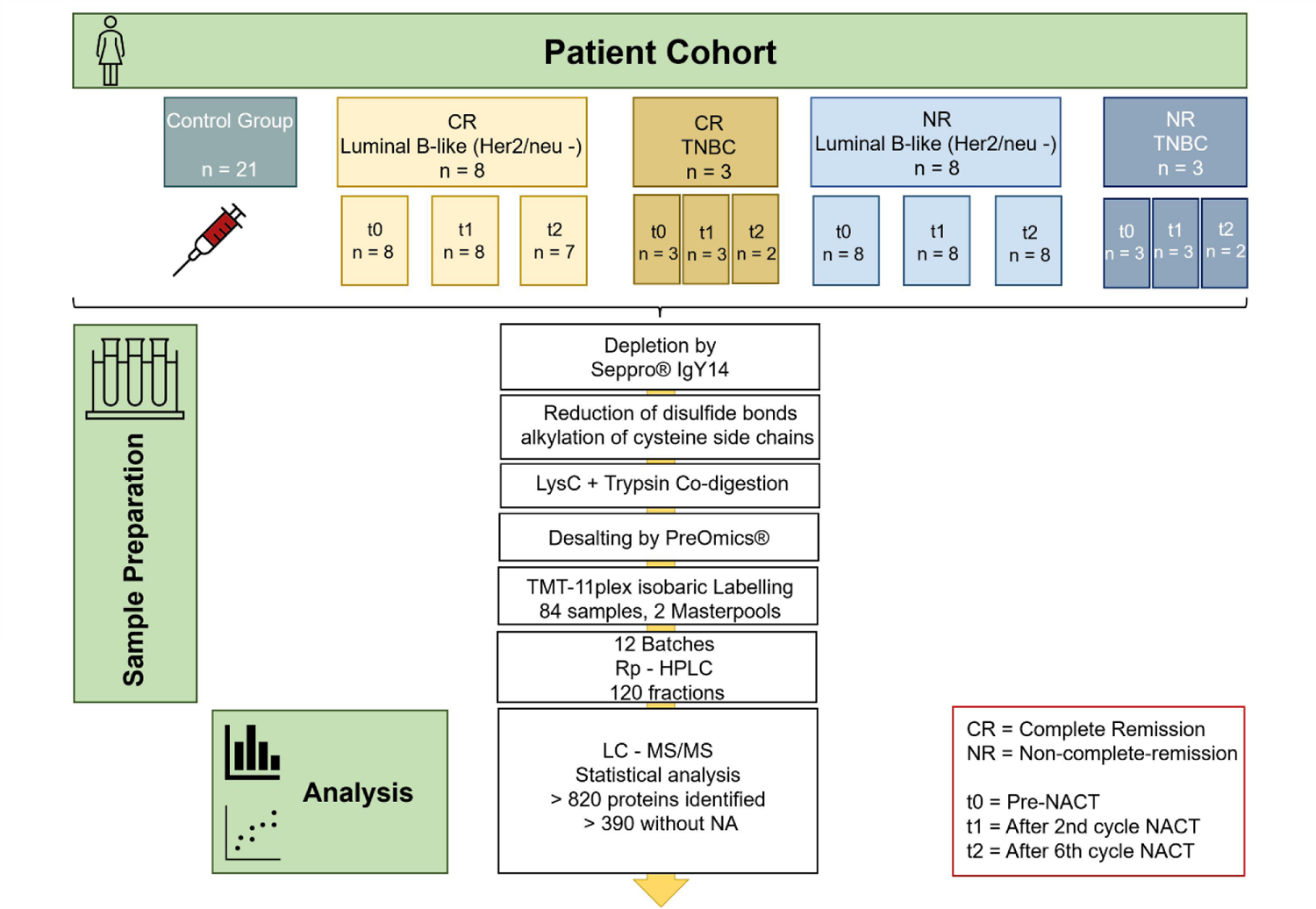
Overview of the proteomic workflow. Patient cohort consisting of 22 patients, 11 patients with pCR (CR) and 11 patients with no pCR (NR). Sera were collected at 3 different time points. Including a control group of 21 healthy female volunteers, 84 serum samples were processed by depletion, digestion, desalting, TMT labelling, fractionation by liquid chromatography and mass spectrometric measurement. TNBC triple negative breast cancer, Rp-HPLC reversed-phase high-performance liquid chromatography, LC - MS/MS liquid chromatography tandem mass spectrometry.

Protein concentration was estimated by bicinchoninic acid (BCA) assay (ThermoScientific). Approximately 580 µg of protein was further processed according to the manufacturer’s protocol of the Seppro® IgY14 immunoaffinity kit. After depletion, the protein concentration was measured again using the BCA assay. Prior to digestion, 50 mM HEPES pH 8.0 (Sigma-Aldrich) was added to the sample to buffer pH interactions with the dilution buffer. 0.1% RapiGest was added, and the sample was placed on a heating plate at 500 rpm for 10 min at 95 °C. To reduce disulphide bonds of cysteine side chains by carbamidomethylation, 2 mM tris(2-carboxyethyl)phosphine (TCEP, Sigma-Aldrich) was added and the sample was placed on a heating plate at 500 rpm for 10 min at 37 °C. Iodoacetamide (IAA, Sigma-Aldrich) was prepared in the dark and 20 mM was added to the sample to alkylate cysteines and prevent disulfide bonds. The sample was placed on a heating plate for 10 min at 500 rpm at 37°C in the dark. To quench excessive IAA, 5 mM dithiothreitol (DTT, Sigma-Aldrich) was added and placed on a heating plate for 10 min at 500 rpm at RT. Lys-C-trypsin co-digestion was performed by adding Lys-C (Wako Chemicals MS grade) at a ratio of 1:100 to the total protein mass in the sample. The sample was covered with parafilm and placed on an end-to-end rotator in an oven at 50°C for 2 h. Trypsin (Promega MS grade) was then added at a ratio of 1:50 to the total protein mass in the sample, covered with parafilm and placed on a heating plate at 37°C overnight (approximately 16 h). To remove RapiGest, a final concentration of 2% trifluoracetic acid (TFA, Sigma-Aldrich) was added to the sample and the sample was placed on a heating plate at 500 rpm for 30 min at 37°C. Peptide clean-up was performed on mixed-phase columns (PreOmics®) according to the manufacturer’s protocol using triethylamine (TEA, Sigma-Aldrich) instead of NH4OH in the elution buffer due to subsequent TMT labelling. Samples were vacuum dried and stored at ™80°C. For further processing, 80 µl of 0.1 M HEPES pH 7.5 was added to each vacuum dried sample and a BCA assay was performed to determine peptide concentration. Randomisation was removed and samples were mass tagged using 10 channels of a TMT-11-plex stock (Thermo Fisher Scientific). For quantification control, different amounts of 11 synthetic peptides (iRT mix) were pipetted onto the samples per batch (2-8µl). Two master pools were generated from all samples. 8 µl TMT-DMSO of a specific channel was added to each sample and incubated at room temperature for 3 h at 600 rpm. Samples from one batch, including both master pools, were pooled in a new 2 ml low-bind tube and stored at ™80 °C. 66 µg of each TMT-labelled pool was fractionated on an Agilent 1100 reversed phase chromatography system. A X Bridge C18 column (150 mm × 1 mm column, 3.5 µm particles) was used, and an increasing linear gradient of acetonitrile (ACN) was applied at a flow rate of 42 µl/min. 36 fractions per batch were collected and combined into 10 fractions (combining fraction 5 with fraction 15 and 25, fraction 6 with fraction 16 and 26 and so forth). Again, iRT peptides were spiked in, followed by vacuum-concentration until dryness. Samples were stored at − 80 °C until LC–MS/MS analysis.

### 2.4 LC-MS/MS Data Acquisition and Analysis

For MS measurement, vacuum-dried samples were dissolved, briefly sonicated and transferred to measurement tubes. Samples were analysed on an Orbitrap Q-Exactive plus mass spectrometer (MS) coupled to an Easy nanoLC 1000 (Thermo Scientific) at a flow rate of 300 nl/min. Peptides were separated on the EASY-Spray™ C18 analytical column (length 250 mm) using an increasing gradient of ACN over 90 min. The MS was operating in a data-dependent acquisition mode and the top 10 precursor ions were targeted for fragmentation scans at 35,000 resolution with 1.2 m/z isolation windows.

Protein identification and quantification was performed by implementing MS-raw files in the MaxQuant platform (V 1-6-7-0). Search parameters were set with cysteine carbamidomethylation as the only fixed modification and trypsin as the protease, with a maximum of two missed cleavages due to Lys-C and trypsin co-digestion. The minimum peptide length was set to 7 amino acids, the MS/MS tolerance to 20 ppm and the PSM and protein false discovery rate to 1%. For random identification, reverse sequences were generated in decoy mode. As isobaric labelling was performed by tandem mass tag (TMT-11plex), reporter ion signals were included per sample. Peptide identification was performed using the Human Proteome Data Set, which includes verified uniprot sequences plus iRT peptides (containing 20426 sequences), and only unique peptides were allowed to be identified. Protein groups known as contaminant, reverse or iRT peptides were removed from further analysis.

Statistical analysis was performed using the open-source statistical software R (V4.0.2). The intensities obtained were log-transformed and normalised using the R package MSStatsTMT (V1.4.6).(19). Statistical analysis was performed on proteins quantified in all patient samples only, resulting in 398 proteins in all 84 samples. Protein significance analyses were performed using the limma package (V.3.44.3) (20). Pairwise comparisons of cycle 2 vs pre-NACT and cycle 6 vs pre-NACT also included a random effect term for intra-patient variability. Unless otherwise stated, analyses were performed separately in the CR and NR subgroups. Differences in intensity levels were assessed by fold change (FC). In addition, the issue of multiple testing was addressed by the Benjamini-Hochberg correction, which adjusted the resulting raw P values by controlling for the false discovery rate (FDR). Further analyses were performed using the EnhancedVolcano package (V1.12.0), the mixOmics package (V6.18.1) for partial least squares discriminant analysis (PLS-DA) and the pROC package (V1.18.0) for ROC curve analysis.

### 2.5 Data availability

LC-MS/MS raw data and analysis output files are available at the European Genome-phenome Archive for appropriate research use (https://ega-archive.org/dacs/EGAC00001002656). As patient-centric proteomic data is increasingly regarded as sensitive, personal data (PMID: 33711481) EGA requires adherence to a data access agreement. The data access agreement for this dataset corresponds to the “Harmonised Data Access Agreement (hDAA) for Controlled Access Data” as brought forward by the “European standardization framework for data integration and data-driven in silico models for personalized medicine – EU-STANDS4PM”.

## 3. Results and Discussion

### 3.1 Patient Cohort and NACT Treatment Scheme

A total of 22 patients were included in the study, of which 11 were in complete remission (pCR) and 11 were not in complete remission (non-pCR). Serum was collected prior to NACT (pre-NACT) and after the second and sixth NACT cycles. The study design strived to enable longitudinal monitoring on a patient-specific basis. Inclusion criteria for this study were a histopathologically confirmed diagnosis of breast cancer, female sex, age over 18 years, planned neoadjuvant chemotherapy and signed informed consent. Exclusion criteria were age under 18 years, post-chemotherapy status, current pregnancy, presence of autoimmune disease, renal insufficiency, HIV infection, acute or chronic hepatitis, participation in a pharmacological study within the previous 3 months, diabetes mellitus, acute infection or history of carcinoma. In addition, 21 sera from healthy female volunteers with no history of carcinoma were included as a control group. Patients were aged between 30 and 66 years (median age 50.3 years) with a primary diagnosis of BC between 2017 and 2019. Healthy volunteers were aged between 24 and 83 years (median age 55.5 years).

All patients were enrolled and treated at the Department of Obstetrics and Gynaecology, University Hospital, University of Freiburg. NACT was administered as a combination of anthracycline (epirubicin E, 90 mg/m2) and alkylating agent (cyclophosphamide C, 600 mg/m2) followed by a taxane derivative (paclitaxel P, 80 mg/m2). Patients received NACT with either a schedule of 4 x EC q3w followed by 4 x P d1, d8, d15 q3w or 4 x EC q2w followed by 4 x P d1, d8, d15 q3w. One patient received only 4 x E q3w followed by 4 x P d1, d8, d15 q3w. After completion of NACT, patients underwent surgery and histopathological examination of the resected tissue. Serum samples were collected at three different time points: pre-NACT (t0), after the second cycle of NACT (after the second cycle of EC) (t1), after the sixth cycle of NACT (after the second cycle of P) (t2) (Figure 2). Additional information, such as grade of differentiation or TNM classification after surgery, is shown in Table I. A total of 84 serum samples were processed and analysed. The cohort and sample size are within the range of recently published serum proteomic studies (21).

**Figure 2.**
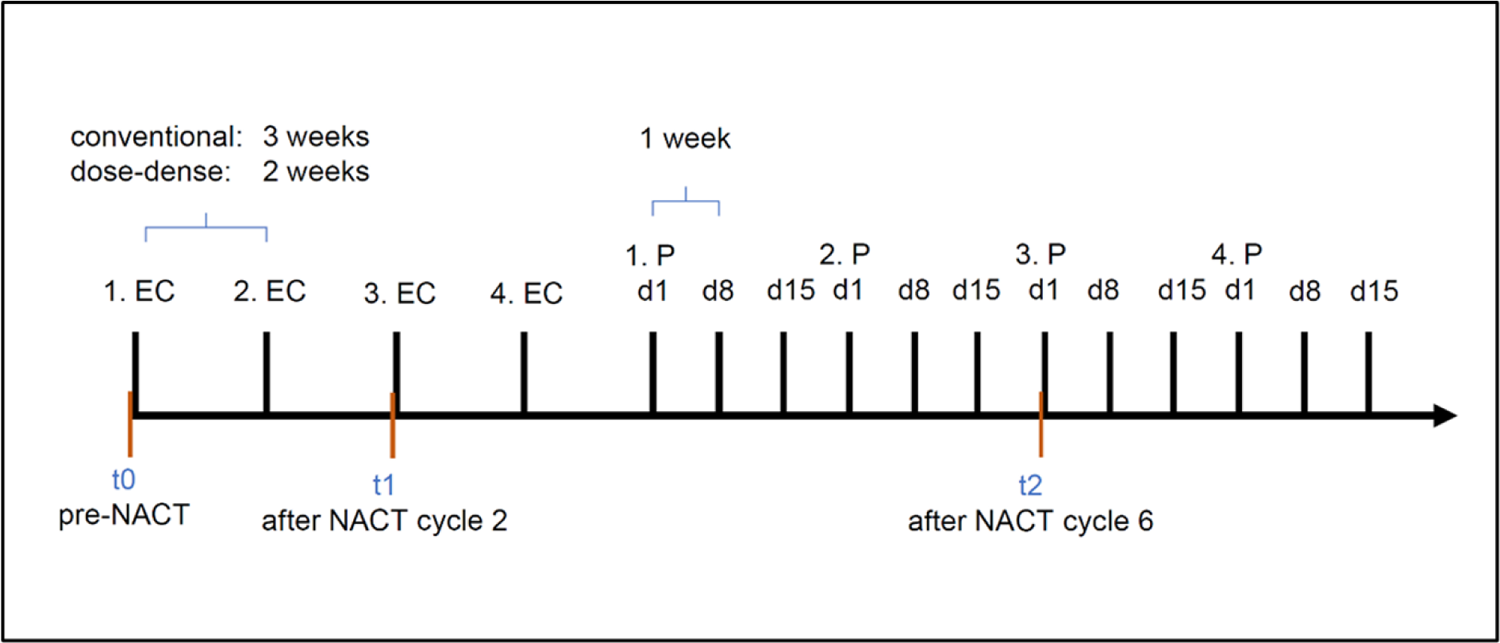
Timeline of collected samples at different timepoints. Samples were collected at time point t0 = pre-NACT, t1 = after cycle 2 of NACT, and t2 = after cycle 6 of NACT. EC Epirubicin Cyclophosphamide, P Paclitaxel

### 3.2 Proteome Coverage

We used a proteomic workflow consisting of high-abundance protein depletion with Seppro® IgY14, TMT-11plex labelling, fractionation and analysis by LC-MS/MS (Figure 1). In a previous study (22), we have successfully benchmarked TMT-based quantitation for a hybrid quadrupole-orbitrap mass spectrometer. The benchmarking also showed an approximately 2-fold ratio compression, meaning that real changes in protein abundance are typically twice as large as indicated by the corresponding TMT-based values. A total of 824 proteins were identified by LC-MS/MS at 1% FDR. Of these, 398 proteins were identified and quantified in each sample (n = 84), representing a “core proteome” free of missingness (Supplement Table I). All further analyses were based on this core proteome. The proteome coverage of 398 proteins is within the range of current serum proteomic studies (21,23–25), although modern fast scanning mass spectrometers reduce the need for depletion of high abundance proteins. The “core proteome” of 398 proteins for the present study spans several orders of quantified protein intensity with a comparable range for all 84 samples (Figure 3), highlighting the successful merging of the 10 TMT label pools used with reference label channels and the corresponding MSStatsTMT algorithms.

**Figure 3.**
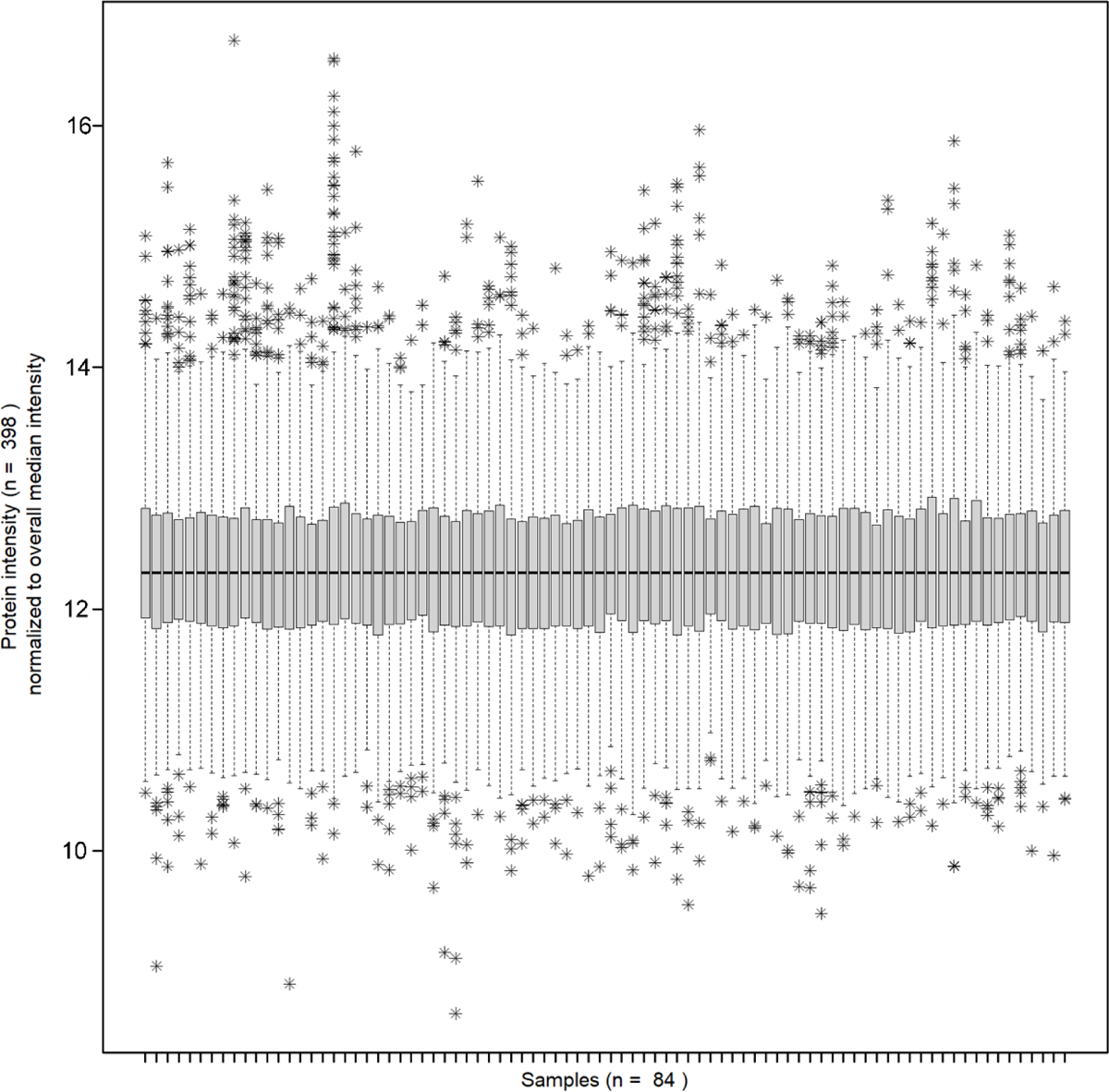
Boxplot depicting the median-normalized protein intensities for 84 serum samples of the present cohort

### 3.3 Global Distinction of Disease Serum Proteomes by Partial Least Squares Discriminant Analysis

To characterise the (dis)similarity of our different conditions, we applied partial least squares discriminant analysis (PLS-DA) using the mixOmics package (26). As a global overview of the distinguishability of the serum proteomes of the included time points of NACT and Ctrl, we observe a pronounced overlap of the corresponding proteome profiles, suggesting only subtle changes in the serum proteome (Figure 4).

**Figure 4.**
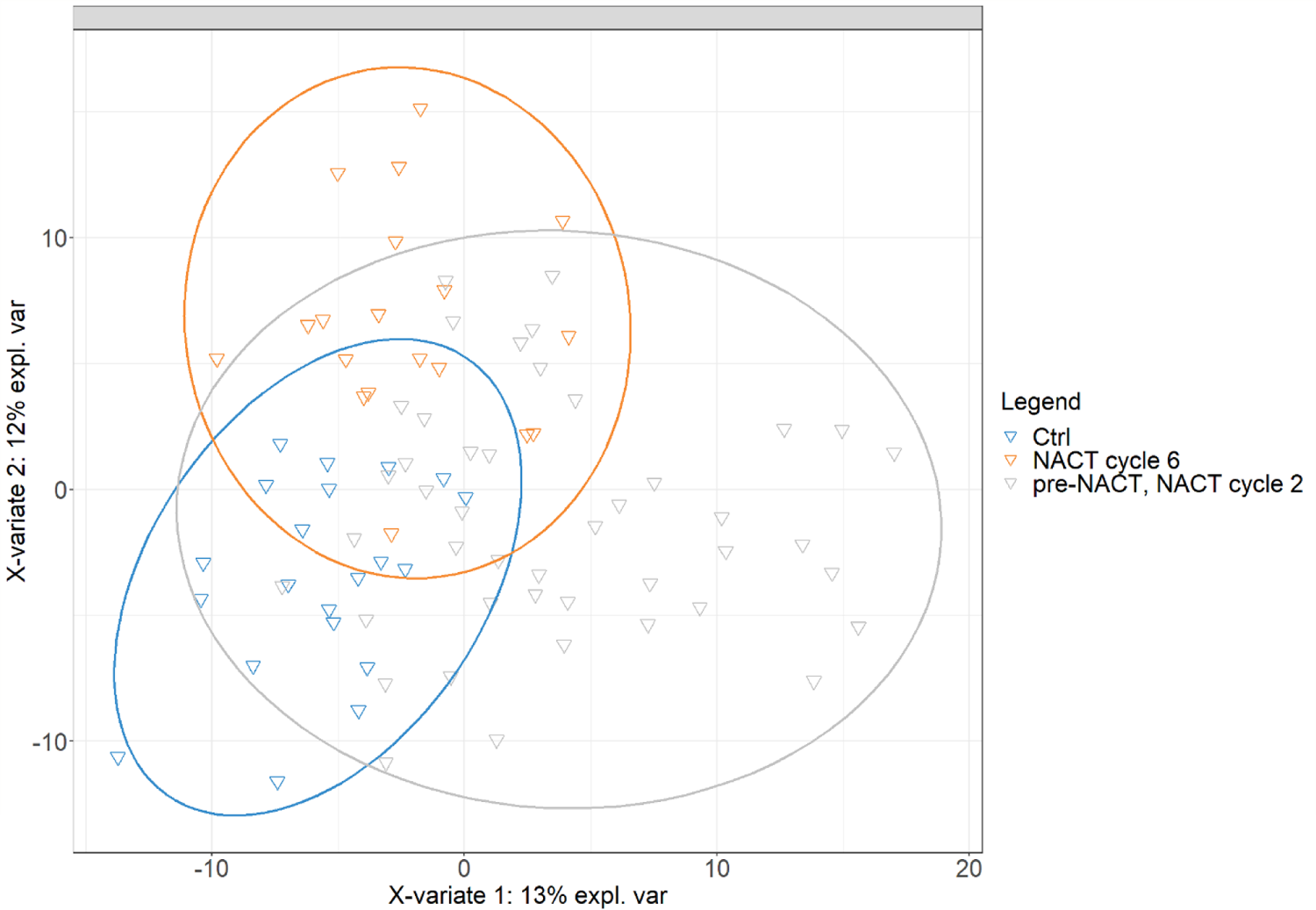
Partial least squares discriminating analysis (PLS-DA). The serum proteomes of the included timepoints during NACT and the Control Group were submitted to PLSDA analysis.

### 3.4 Serum proteome alterations after the second cycle of NACT

For serum proteome comparisons, a pairwise multigroup LIMMA approach was used by creating comparisons between conditions “CR_t1 vs CR_t0, NR_t1 vs NR_t0, CR_t2 vs CR_t0 and NR_t1 vs NR_t0” or “CR_t0 vs Ctrl, NR_t0 vs Ctrl”. For comparisons over time, we implemented a random patient effect in the LIMMA design and used mixed generalised least squares regression. Significantly affected proteins were defined according to the following criteria: (a) FDR-controlled p-value < 0.05, (b) enrichment or depletion of TMT-based protein intensity ≥ 25%. When comparing the serum proteomes of NACT cycle 2 and pre-NACT conditions, several proteins showed significant changes in TMT-based protein intensity. The corresponding volcano plots are shown in Figure 5. For the CR group, 10 proteins met our criteria (Table II). The proteome changes for the NR group were less pronounced, with only 2 proteins meeting the criteria (Table III).

**Figure 5.**
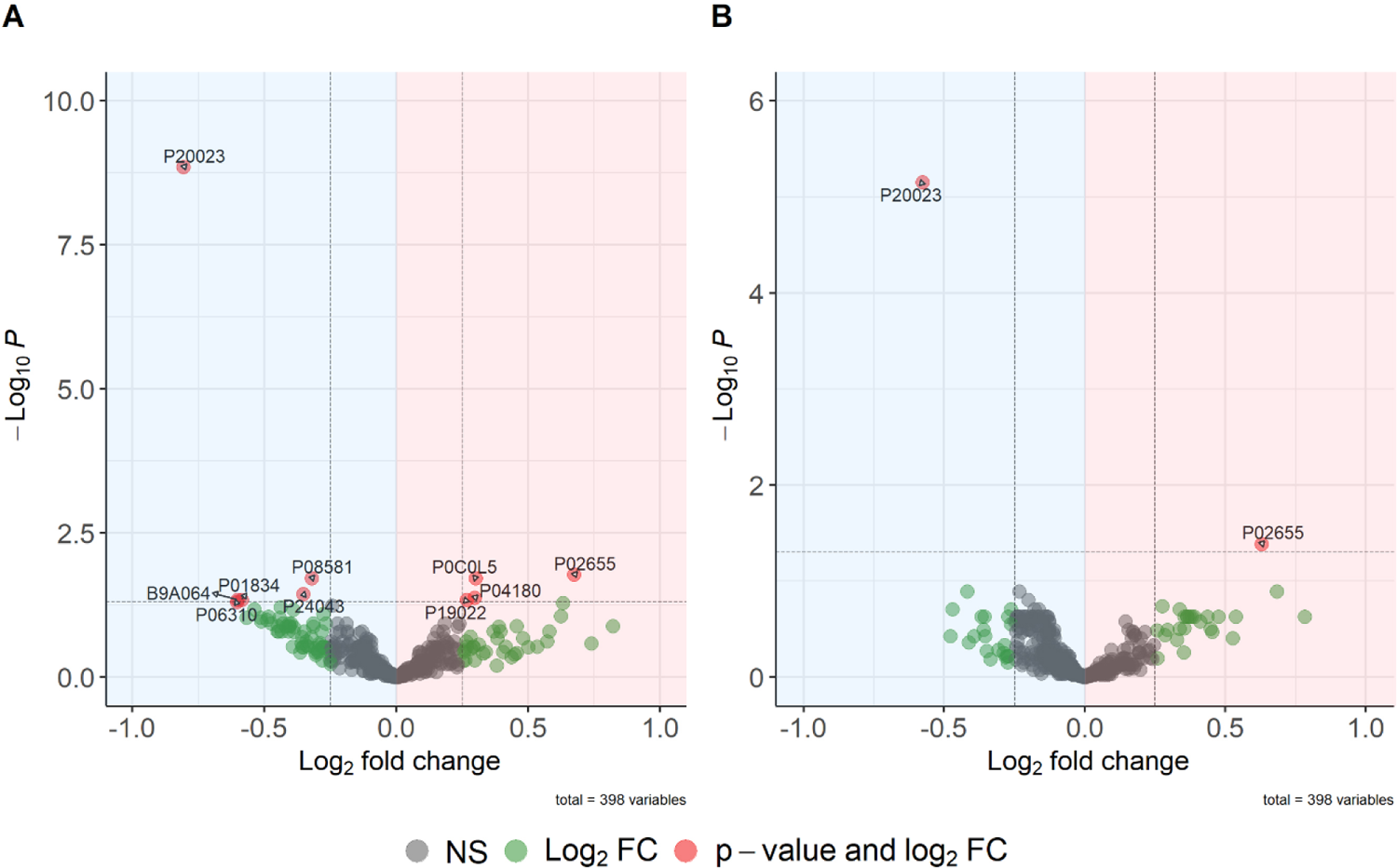
Volcano plots showing the differential protein abundance by pairwise comparisons in limma (“CR_t1 vs. CR_t0” A, “NR_t1 vs. NR_t0” B). Protein intensities after NACT cycle 2 compared to pre-NACT intensities are shown for CR (A) and NR (B). The log2 fold changes (log2FC) are plotted on the x-axis and the corresponding adjusted p-values in -log10 scale are plotted on the y-axis. Significance was defined by an adjusted p-value < 0.05, log2FC ≥ 0.25 (proteins meeting these criteria are coloured red and marked with their corresponding UniProt ID). Upregulated proteins (log2FC > 0) are highlighted in red, downregulated proteins (log2FC < 0) are highlighted in blue. Fold change is expected to be underestimated by 2-fold ratio compression due to TMT labelling.

**Table II.** Significantly affected proteins after the second cycle of NACT in the group of CR compared to pre-NACT baseline.

| Protein name | UniProt ID | Adj. p-value | Log2(t1/t0) |
| --- | --- | --- | --- |
| CR2 | P20023 | < 0.000001 | -0.81 |
| c-MET / HGF receptor | P08581 | 0.02 | -0.38 |
| Immunoglobulin kappa constant | P01834 | 0.04 | -0.59 |
| Immunoglobulin kappa variable 2-30 | P06310 | 0.05 | -0.60 |
| Immunoglobulin lambda-like polypeptide 5 | B9A064 | 0.04 | -0.60 |
| Laminin subunit alpha-2 | P24043 | 0.04 | -0.35 |
| Apolipoprotein C-II | P02655 | 0.02 | 0.67 |
| Basic complement C4 | P0C0L5 | 0.02 | 0.30 |
| N-Cadherin | P19022 | 0.04 | 0.27 |
| Phosphatidylcholine-sterol acyltransferase | P04180 | 0.04 | 0.30 |

Proteins listed with significant differential abundances by pairwise comparisons in limma ("CR\_t1 vs CR\_t0"). Proteins are listed by their names with their corresponding UniProt ID, adjusted p-value and fold change (Binary logarithm of intensity\_t1/intensity\_t0), which is expected to be underestimated by 2-fold Ratio-Compression due to TMT-Labeling.

**Table III.** Significantly affected proteins after the second cycle of NACT in the group of NR compared to pre-NACT baseline.

| Protein name | UniProt ID | Adj. p-value | log2(t1/t0) |
| --- | --- | --- | --- |
| Complement receptor 2 | P20023 | < 0.00001 | -0.56 |
| Apolipoprotein C-II | P02655 | 0.04 | 0.63 |

Proteins listed with significant differential abundances by pairwise comparisons in limma ("NR\_t1 vs NR\_t0"). Proteins are listed by their names with their corresponding UniProt ID, adjusted p-value and fold change (Binary logarithm of intensity\_t1/intensity\_t0), which is expected to be underestimated by 2-fold Ratio-Compression due to TMT-Labeling.

For the protein c-Met (P08581), we observed significantly decreased levels (adjusted p-value = 0.02) in the CR group after the second cycle of NACT compared to pre-NACT baseline, which normalised to pre-NACT baseline after the third cycle (Figure 6, A). C-Met levels did not appear to be affected in the non-responder group, nor did the intensity levels over time differ from the control group (Figure 6, B). C-Met is encoded by the MET gene and functions as a transmembrane receptor tyrosine kinase that binds hepatocyte growth factor (HGF) and is predominantly found on epithelial and endothelial cells. Amplification and polysomy of MET, as well as protein overexpression of c-Met, have previously been described in breast cancer tissue and may be associated with poorer prognosis (27,28). With carbozatinib, c-Met is already an established therapeutic target. As an inhibitor of the tyrosine kinases c-Met and VEGFR2, the c-Met/HGF signalling pathway is downregulated in medullary thyroid cancer, renal cell carcinoma and hepatocellular carcinoma. Current phase II studies show significant benefit of carbozatinib therapy in metastatic breast cancer (29–34).

**Figure 6.**
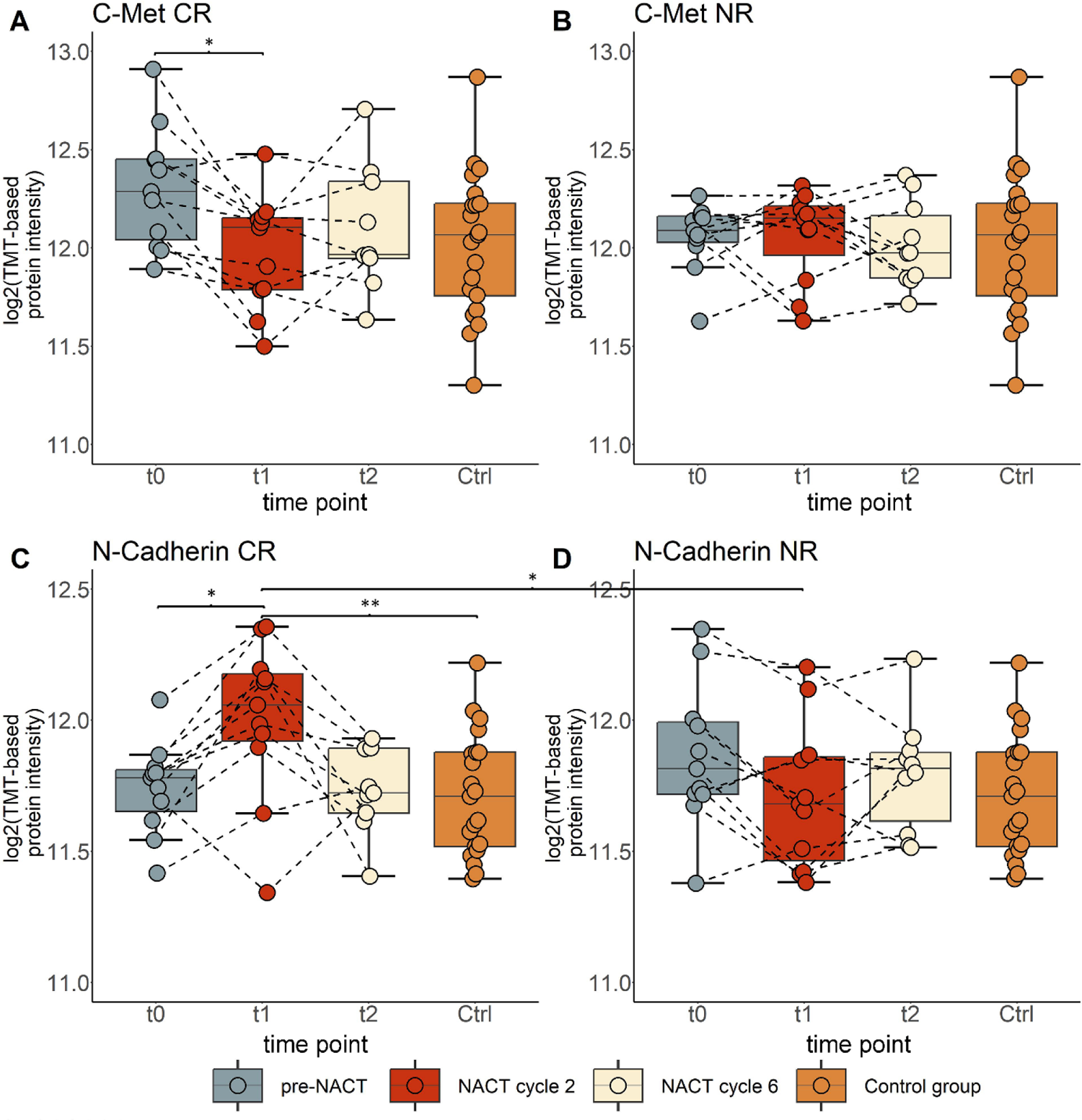
Boxplots illustrating the serum abundance of C-Met (A, B) and N-cadherin (C, D) in CR and NR over time (before NACT, after NACT cycle 2, after NACT cycle 6) and in the control group. Abundances are expressed as log2 TMT-based protein intensities. Differential protein abundance analysis was performed by pairwise comparisons in Limma. Significance was defined by an adjusted p-value < 0.05 and highlighted as follows: *adjusted p-value < 0.05, **adjusted p-value < 0.01. Samples from one patient are connected by a line.

For N-cadherin (P19022), we observed significantly increased intensity levels in the CR group after the second cycle of NACT compared to pre-NACT baseline (adjusted p-value = 0.04), which normalised to pre-NACT baseline after the sixth cycle (Figure 6, C). N-cadherin levels did not appear to be affected in the non-responder group, nor did pre-NACT levels differ from the control group (Figure 6, D). However, we reported significantly dysregulated levels of N-cadherin comparing CR after the second cycle of NACT to Ctrl (adjusted p-value < 0.01, Figure 6, C) and comparing the CR group to NR after the second cycle of NACT (adjusted p-value = 0.03) (Figure 6, C-D). In addition to P-, R- and M-cadherin, E- and N-cadherin belong to the subgroup of classical type I cadherins. E-cadherin is located on the cell membrane of epithelial cells and is responsible for strong cell-cell contacts. N-cadherin, found on non-epithelial tissues, modulates cell differentiation during cell development and plays a central role in angiogenesis by stabilising endothelial cells and pericytes. N-cadherin has been reported to be upregulated in several cancers, whereas E-cadherin is often downregulated (35,36). Furthermore, according to ProteinAtlas.org, N-cadherin is one of the 300-500 most abundant serum proteins. The results of our study show significantly increased levels of N-cadherin after the second cycle of NACT. As we defined CR as patients with pathological complete remission, N-cadherin has the potential to be a predictive marker for pCR. Furthermore, due to the inverse abundance of C-Met and N-cadherin over time, we see predictive potential in an N-cadherin/c-Met ratio. The time-dependent delta (intensity_t1 - intensity_t0) of c-Met and N-cadherin showed a high predictive value (AUC = 0.93) in classifying patients in terms of their response (either NR or CR) (Figure 7).

**Figure 7.**
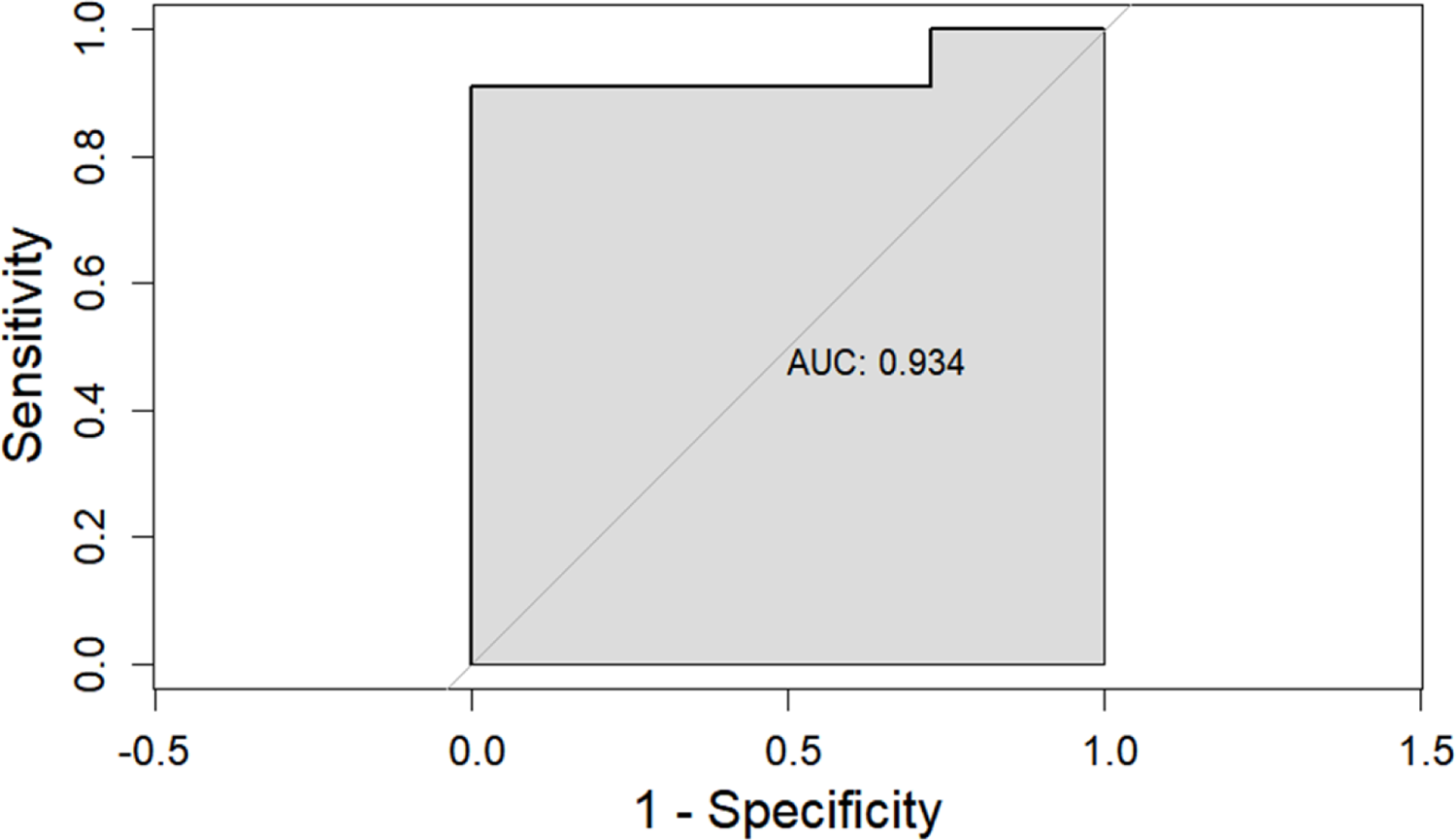
Receiver operating characteristic (ROC) curve for time dependent delta of c-Met and N-Cadherin as protein classifier.

### 3.5 Serum proteome alterations after the sixth cycle of NACT

After the sixth cycle of NACT (after the second cycle of P), we observed a markedly higher number of affected serum proteins in both groups compared to the second cycle of NACT. In the CR group, 23 proteins showed significantly dysregulated intensity levels, of which 13 proteins were downregulated and 10 proteins were upregulated (Figure 8, A, Table IV). In the NR group, we observed a significantly higher number of significantly dysregulated proteins (#38) compared to CR (Figure 8, B, Table V).

**Figure 8.**
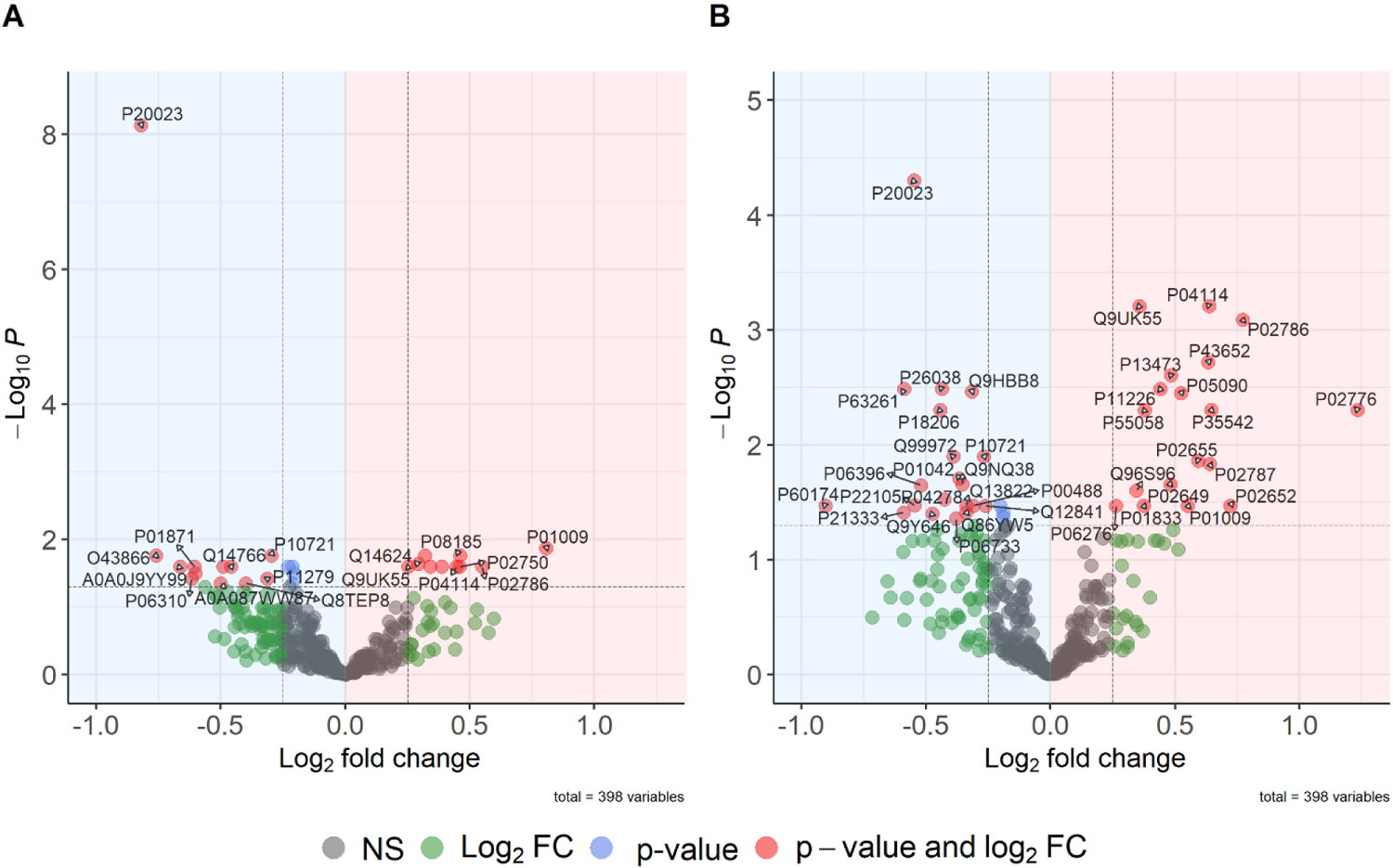
Volcano plots showing the differential protein abundance by pairwise comparisons in limma (“CR_t2 vs. CR_t0” A, “NR_t2 vs. NR_t0” B). Protein intensities after NACT cycle 6 compared to pre-NACT intensities are shown for CR (A) and NR (B). The log2 fold changes (log2FC) are plotted on the x-axis and the corresponding adjusted p-values in - log10 scale are plotted on the y-axis. Significance was defined by an adjusted p-value < 0.05, log2FC ≥ 0.25 (proteins meeting these criteria are coloured red and marked with their corresponding UniProt ID). Upregulated proteins (log2FC > 0) are highlighted in red, downregulated proteins (log2FC <0) are highlighted in blue. Fold change is expected to be underestimated by 2-fold ratio compression due to TMT labelling.

**Table IV.**
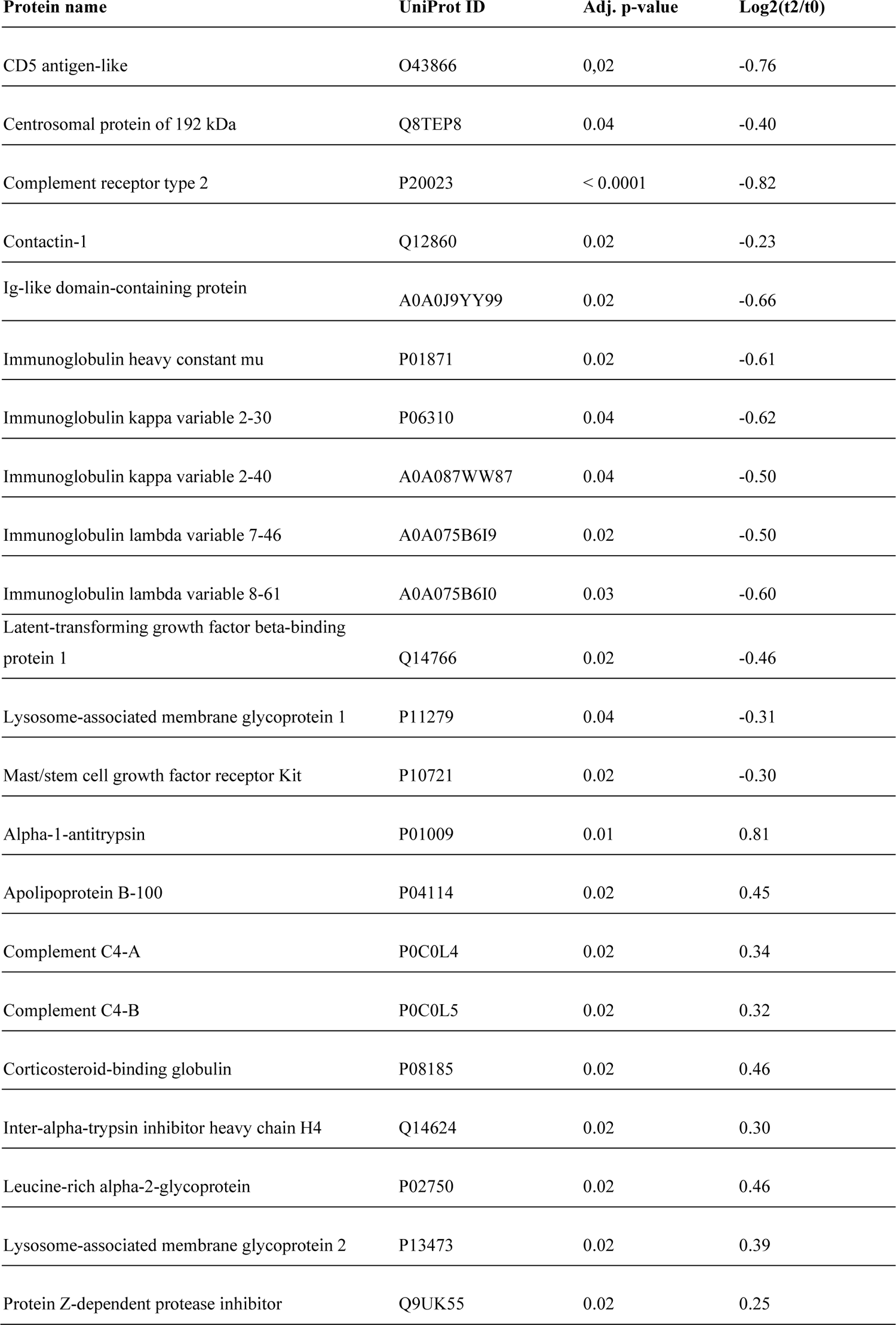
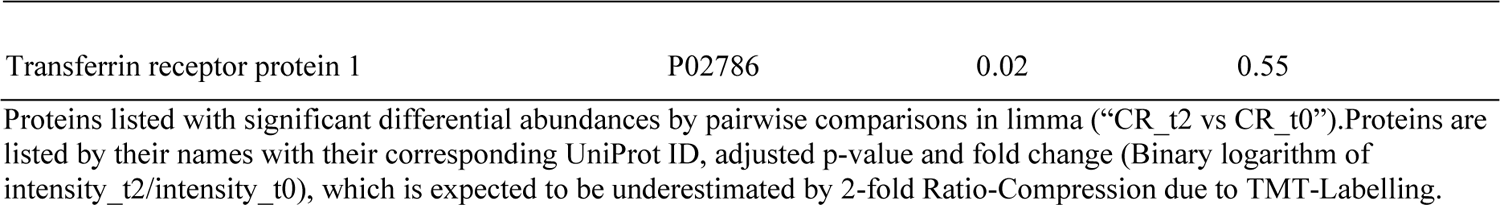
Significantly affected proteins after the sixth cycle of NACT in CR compared to pre-NACT baseline.

**Table V.** Significantly affected proteins after the sixth cycle of NACT in NR compared to pre-NACT baseline.

| Protein name | UniProt ID | Adj. p-value | Log2(t2/t0) |
| --- | --- | --- | --- |
| Actin, cytoplasmic 2 | P63261 | < 0.01 | -0.59 |
| Alpha-enolase | P06733 | 0.04 | -0.38 |
| Cadherin-related family member 5 | Q9HBB8 | < 0.01 | -0.32 |
| Carboxypeptidase Q | Q9Y646 | 0.04 | -0.47 |
| Coagulation factor XIII A chain | P00488 | 0.03 | -0.31 |
| Complement receptor type 2 | P20023 | < 0.0001 | -0.55 |
| Ectonucleotide pyrophosphatase/phosphodiesterase family member 2 | Q13822 | 0.03 | -0.34 |
| Filamin-A | P21333 | 0.04 | -0.59 |
| Follistatin-related protein 1 | Q12841 | 0.03 | -0.26 |
| Gelsolin | P06396 | 0.02 | -0.52 |
| Kininogen-1 | P01042 | 0.02 | -0.37 |
| Mast/stem cell growth factor receptor Kit | P10721 | 0.01 | -0.27 |
| Moesin | P26038 | < 0.01 | -0.44 |
| Myocilin | Q99972 | 0.01 | -0.39 |
| Serine protease inhibitor Kazal-type 5 | Q9NQ38 | 0.02 | -0.35 |
| Sex hormone-binding globulin | P04278 | 0.03 | -0.43 |
| Tenascin-X | P22105 | 0.03 | -0.54 |
| Trem-like transcript 1 protein | Q86YW5 | 0.04 | -0.34 |
| Triosephosphate isomerase | P60174 | 0.03 | -0.90 |
| Vinculin | P18206 | < 0.01 | -0.44 |
| Afamin | P43652 | < 0.01 | 0.63 |
| Alpha-1-antitrypsin | P01009 | 0.03 | 0.55 |
| Apolipoprotein A-II | P02652 | 0.03 | 0.72 |
| Apolipoprotein B-100 | P04114 | < 0.001 | 0.64 |
| Apolipoprotein C-II | P02655 | 0.01 | 0.59 |
| Apolipoprotein D | P05090 | < 0.01 | 0.52 |
| Apolipoprotein E | P02649 | 0.02 | 0.48 |
| Cholinesterase | P06276 | 0.03 | 0.26 |
| Lysosome-associated membrane glycoprotein 2 | P13473 | < 0.01 | 0.48 |
| Mannose-binding protein C | P11226 | < 0.01 | 0.44 |
| Phosphatidylethanolamine-binding protein 4 | Q96S96 | 0.02 | 0.34 |
| Phospholipid transfer protein | P55058 | < 0.01 | 0.38 |
| Platelet factor 4 | P02776 | < 0.01 | 1.23 |
| Polymeric immunoglobulin receptor | P01833 | 0.03 | 0.37 |
| Protein Z-dependent protease inhibitor | Q9UK55 | < 0.001 | 0.36 |
| Serotransferrin | P02787 | 0.01 | 0.64 |
| Serum amyloid A-4 protein | P35542 | < 0.01 | 0.64 |
| Transferrin receptor protein 1 | P02786 | < 0.001 | 0.77 |

Proteins listed with significant differential abundances by pairwise comparisons in limma ("NR\_t2 vs NR\_t0"). Proteins are listed by their names with their corresponding UniProt ID, adjusted p-value and fold change (Binary logarithm of intensity\_t2/intensity\_t0), which is expected to be underestimated by 2-fold Ratio-Compression due to TMT-Labeling.

When comparing protein intensity levels over time between CR and NR, four proteins with significantly dysregulated intensities showed inverse behaviour between conditions (centrosomal protein, contactin-1, sex hormone-binding globulin, cholinesterase, Figure 9). The intensity of contactin-1 (Q12860), a neuronal transmembrane glycoprotein, decreased significantly in CR after six cycles of NACT (adjusted p-value = 0.02). Interestingly, it showed a general downregulation of intensity levels in the NR group (Figure 9, A). Contactin-1 is known to play a central role in cell adhesion. Overexpression of contactin-1 has been reported in various cancer entities (37). For the centrosomal protein 192 kDa (Q8TEP8), we measured significantly decreased levels after six cycles of NACT (adjusted p-value = 0.04), whereas its levels remained stable in the NR group (Figure 9, B).

**Figure 9.**
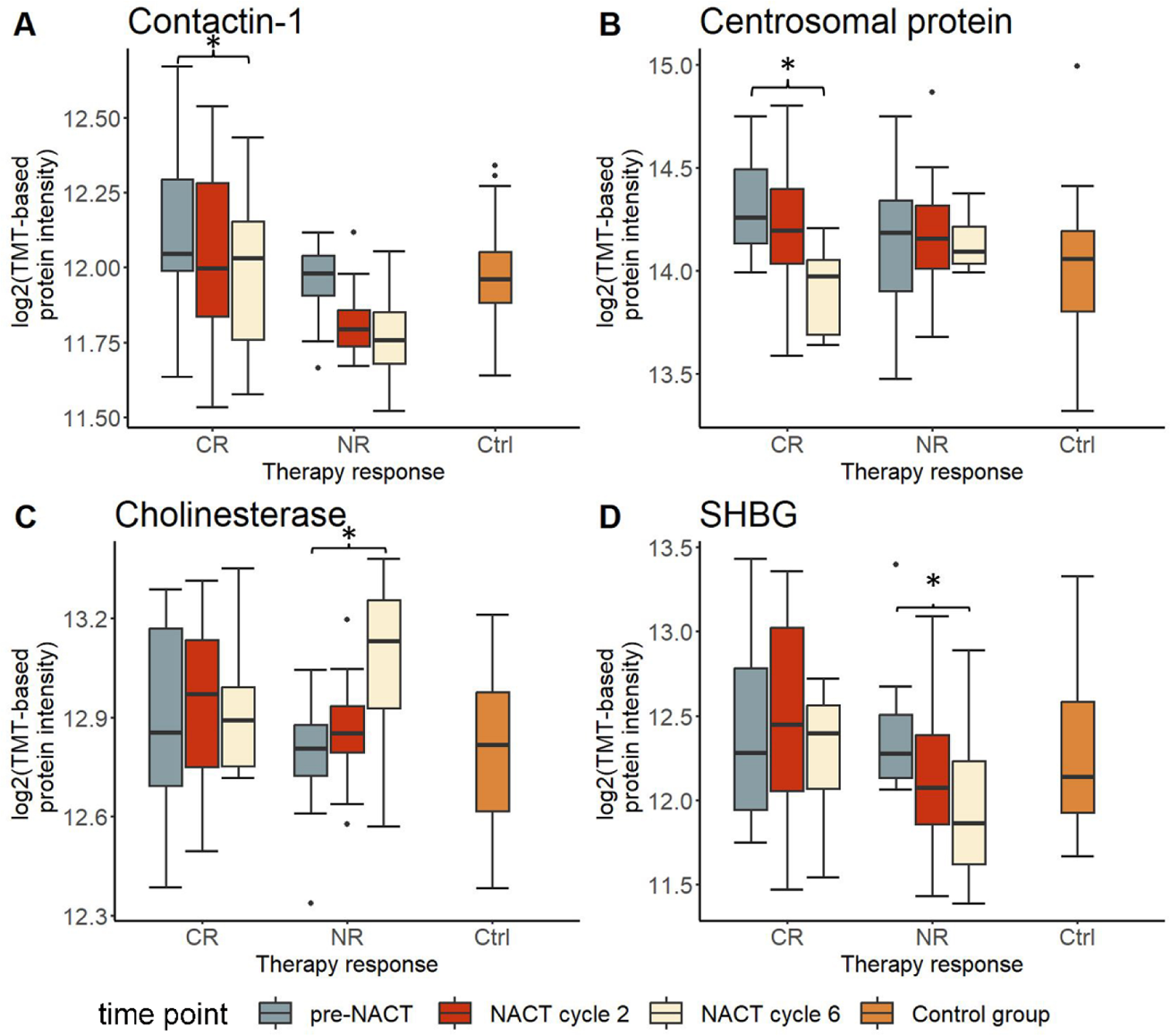
Boxplots illustrating the serum abundance of contactin-1, centrosomal protein, sex hormone-binding globulin and cholinesterase over time (pre-NACT, after NACT cycle 2, after NACT cycle 6) in CR and NR and in the control group. Abundances are expressed as log2 TMT-based protein intensities. Differential protein abundance analysis was performed by pairwise comparisons in Limma. Significance was highlighted as follows: *adjusted p-value < 0.05.

Furthermore, levels of the cholinesterase butyrylcholinesterase (P06276) showed a significant increase in the NR group after six cycles of NACT (adjusted p-value = 0.03), whereas levels remained stable in the CR group (Figure 9, C).

Sex hormone-binding globulin (SHBG, P04278) caught our attention with significantly downregulated levels in the NR group after six cycles of NACT (adjusted p-value = 0.03), while there was no significant change over time in the CR group (Figure 9, D). SHBG is mainly produced in the liver, but also in the brain, uterus, testes, placenta and vagina. Looking more closely at the subtypes included in this study, we observe an inverse behaviour between Luminal B-like (HER2/neu-) and TNBC within the group of NR (Figure 10). Here, an increase in SHBG intensity was measured before NACT in TNBC (Figure 10, C). The intensity decreased after the second cycle of NACT and increased again after the sixth cycle in TNBC. In luminal B-like (Her2/neu) samples, intensities were generally lower and decreased over time in the NR group. However, a robust statistical analysis is lacking due to the small number of TNBC samples.

**Figure 10.**
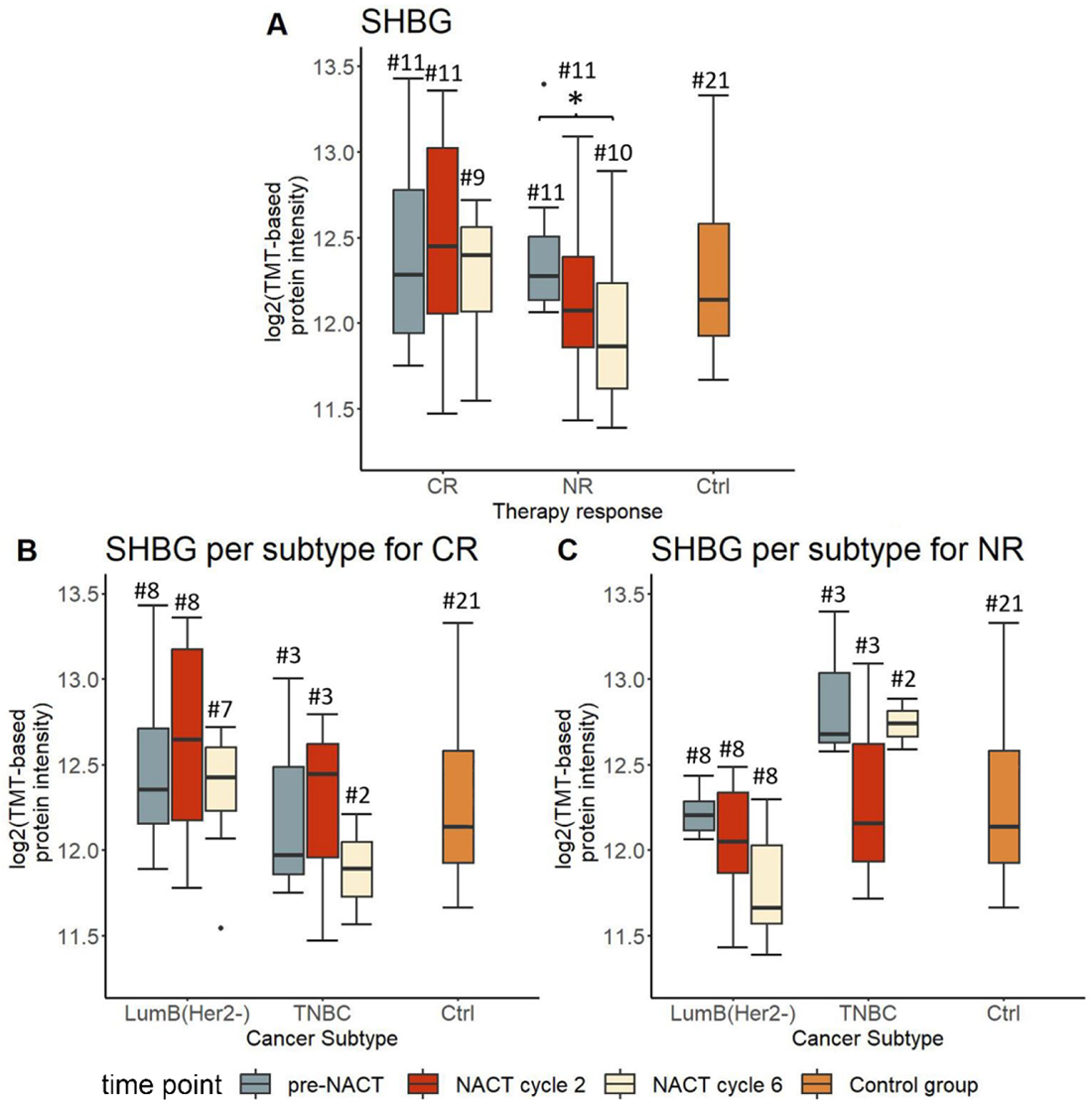
Boxplots illustrating the serum abundance of SHBG over time (pre-NACT, after NACT cycle 2, after NACT cycle 6), per therapy response (A) and per subtype in CR (B) and NR (C). Abundances displayed as log2 TMT-based protein intensities. Differential protein expression analysis was performed by pairwise comparisons in limma. Significance was highlighted as follows: *adjusted p-value < 0.05.

### 3.6 Serum proteome comparisons between complete remission and non-complete remission pre-NACT, after NACT cycle 2 and after NACT cycle 6

No significant differences were measured comparing protein intensity levels of CR to NR before NACT (Figure 11, A). After cycle 2, one protein (N-cadherin) met the criteria for significant changes in intensity by enrichment (Figure 11, B). As described earlier, N-cadherin also showed an inverse behaviour after two cycles of NACT when comparing fold changes between CR and NR. After 6 cycles of NACT, 7 proteins met our criteria for significance, but none showed an inverse behaviour in fold changes comparing CR with NR (Figure 11, C, Table VI).

**Figure 11.**
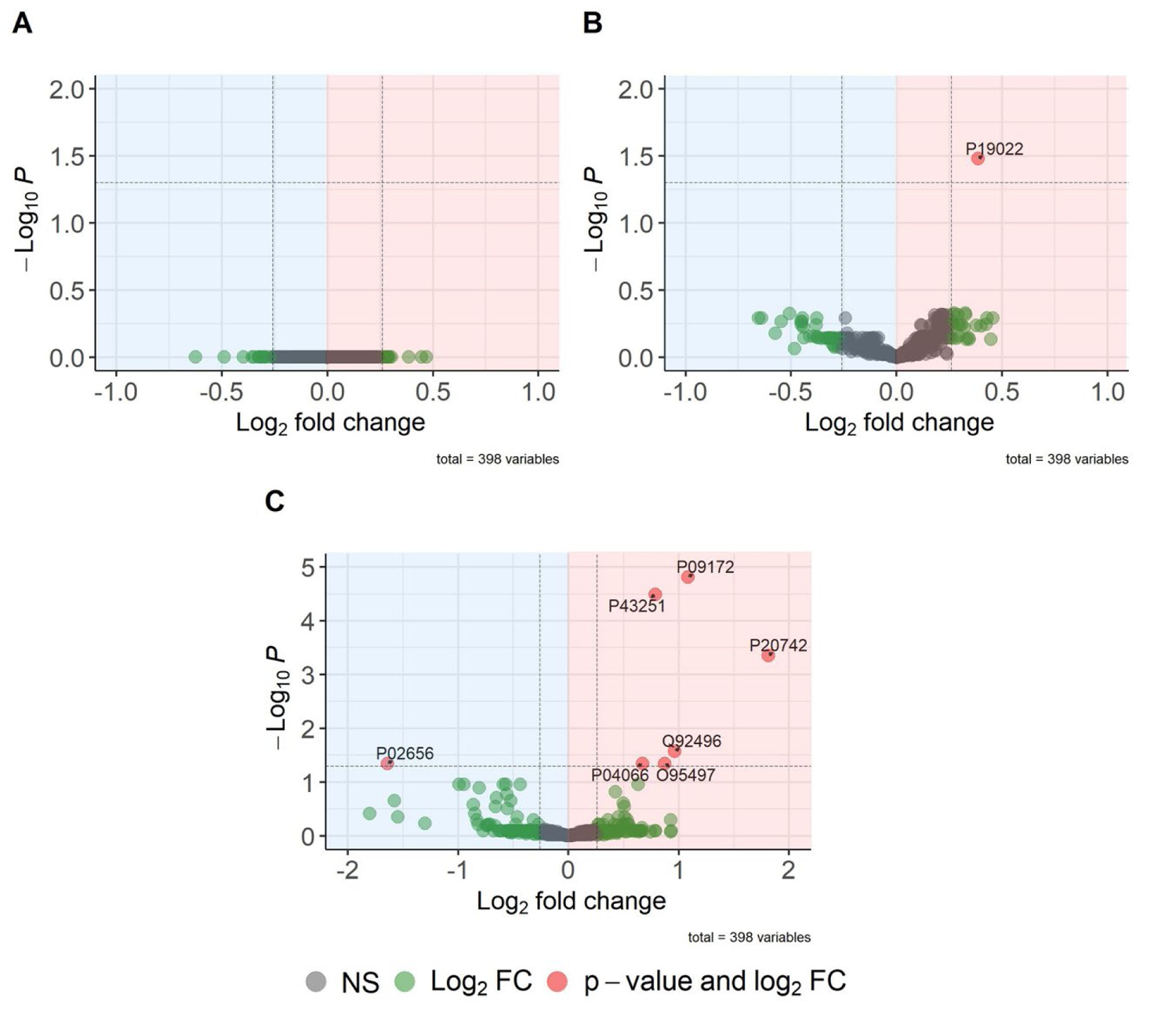
Volcano plots showing the differential protein abundance by pairwise comparison in limma (“CR_t0 vs. NR_t0” A, “CR_t1 vs. NR_t1” B, “CR_t3 vs. NR_t3” C). Protein intensities CR pre-NACT compared to NR pre-NACT (A), CR after NACT cycle 2 compared to NR after NACT cycle 2 (B) and CR after NACT cycle 6 compared to NR after NACT cycle 6 (C) are shown. The log2 fold changes (log2FC) are plotted on the x-axis and the corresponding adjusted p-values in - log10 scale are plotted on the y-axis. Significance was defined by an adjusted p-value < 0.05, log2FC ≥ 0.25.

**Table VI.**
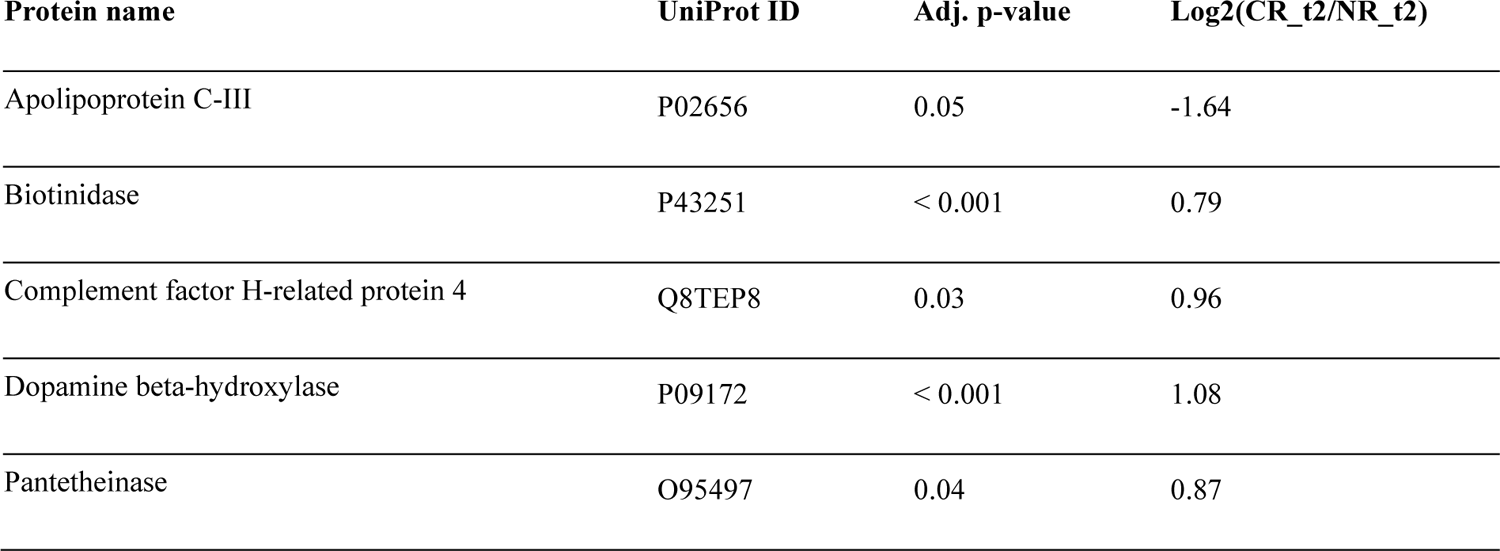
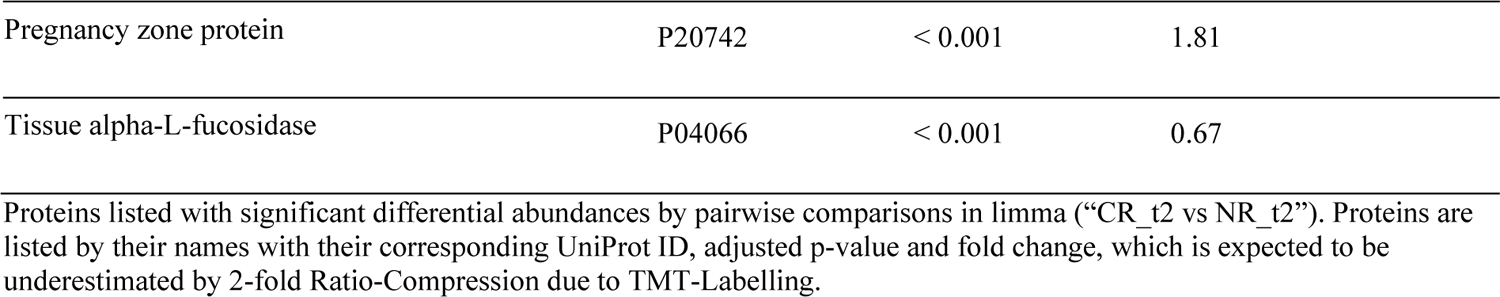
Significantly differential regulated proteins comparing CR with NR after the sixth cycle of NACT.

| Protein name | UniProt ID | Adj. p-value | Log<sub>2</sub>(CR_t2/NR_t2) |
| --- | --- | --- | --- |
| Apolipoprotein C-III | P02656 | 0.05 | -1.64 |
| Biotinidase | P43251 | < 0.001 | 0.79 |
| Complement factor H-related protein 4 | Q8TEP8 | 0.03 | 0.96 |
| Dopamine beta-hydroxylase | P09172 | < 0.001 | 1.08 |
| Pantetheinase | O95497 | 0.04 | 0.87 |
| Pregnancy zone protein | P20742 | < 0.001 | 1.81 |
| Tissue alpha-L-fucosidase | P04066 | < 0.001 | 0.67 |

Proteins listed with significant differential abundances by pairwise comparisons in limma ("CR\_t2 vs NR\_t2"). Proteins are listed by their names with their corresponding UniProt ID, adjusted p-value and fold change, which is expected to be underestimated by 2-fold Ratio-Compression due to TMT-Labeling.

### 3.7 Serum proteome comparison of pre-NACT baseline and control group

In addition, we compared the proteomic profile of the pre-NACT serum samples with the control group consisting of age-matched healthy female volunteers. Here, 61 proteins showed significantly dysregulated intensities, of which 42 proteins were downregulated in BC pre-NACT samples, whereas 19 proteins were enriched (Figure 12, Table VII).

**Figure 12.**
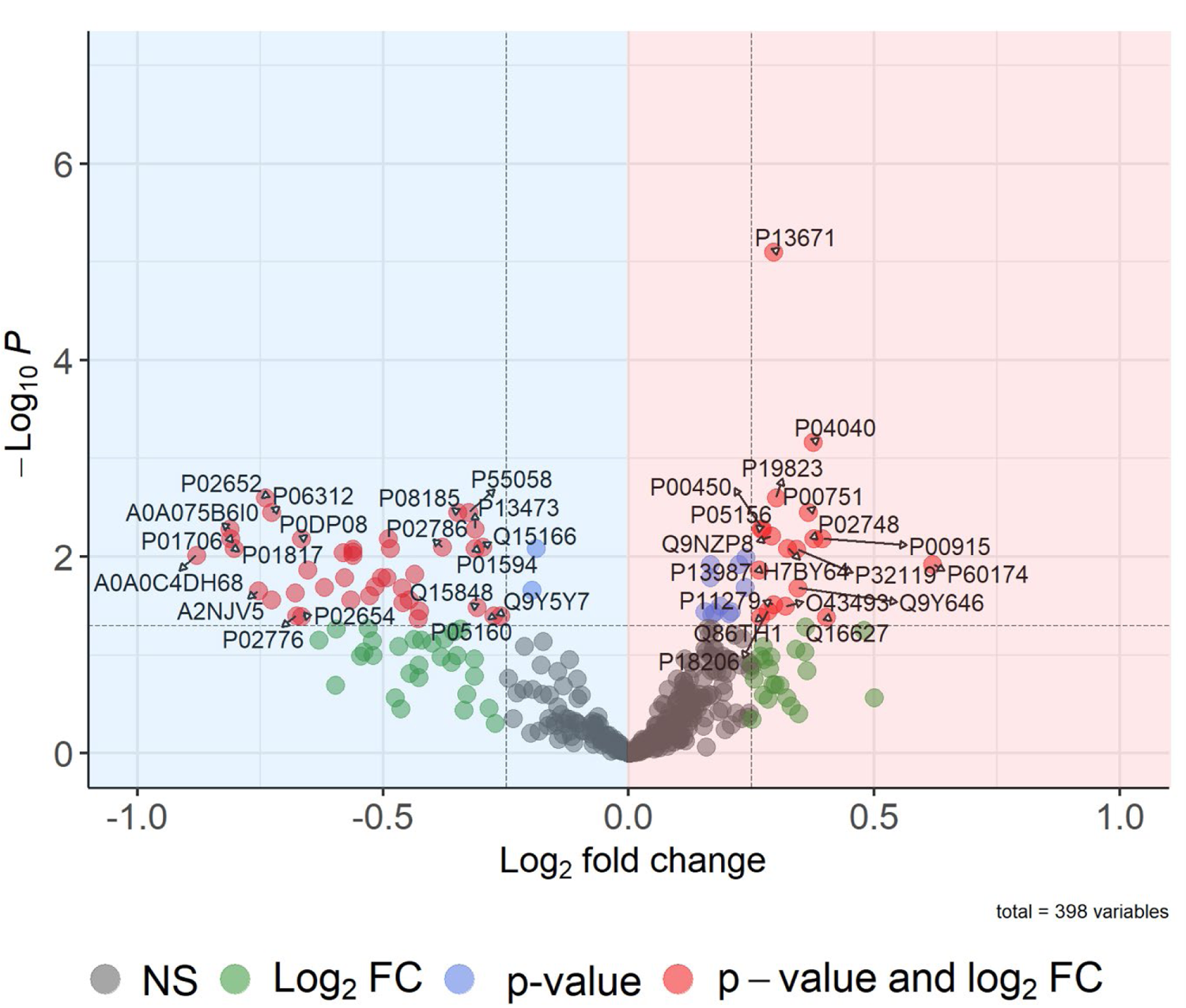
Volcano plot showing the differential protein abundance by pairwise comparison in limma (“CR/NR_t0 vs. Ctrl”). Protein intensities for BC patients before NACT were compared to the control group by pairwise comparison in limma. The log2 fold changes (log2FC) are plotted on the x-axis and the corresponding adjusted p-values in -log10 scale are plotted on the y-axis. Significance was defined by an adjusted p-value < 0.05, log2FC ≥ 0.25 (proteins meeting these criteria are coloured red and marked with their corresponding UniProt ID). Fold change is expected to be underestimated by 2-fold ratio compression due to TMT labelling.

**Table VII.** Significantly differential regulated proteins in BC patients pre-NACT compared to the Control Group.

| Protein name | UniProt ID | Adj. p-value | Log2(BC_t0/Ctrl) |
| --- | --- | --- | --- |
| Adiponectin | Q15848 | 0.02 | -0.33 |
| Alpha-2-macroglobulin | P01023 | 0.02 | -0.56 |
| Apolipoprotein A-I | P02647 | 0.01 | -0.48 |
| Apolipoprotein A-II | P02652 | < 0.01 | -0.74 |
| Apolipoprotein C-I | P02654 | 0.03 | -0.68 |
| Coagulation factor XIII B chain | P05160 | 0.03 | -0.30 |
| Corticosteroid-binding globulin | P08185 | < 0.01 | -0.33 |
| Ig-like domain-containing protein | A0A0J9YY99 | 0.01 | -0.50 |
| Immunoglobulin heavy constant mu | P01871 | 0.04 | -0.62 |
| Immunoglobulin heavy variable 1-69 | P01742 | 0.01 | -0.53 |
| Immunoglobulin heavy variable 2-5 | P01817 | 0.01 | -0.81 |
| Immunoglobulin heavy variable 3/OR16-9 | S4R460 | 0.04 | -0.36 |
| Immunoglobulin heavy variable 3-15 | A0A0B4J1V0 | 0.01 | -0.63 |
| Immunoglobulin heavy variable 3-43D | P0DP04 | 0.02 | -0.46 |
| Immunoglobulin heavy variable 3-64D | A0A0J9YX35 | 0.05 | 0.41 |
| Immunoglobulin heavy variable 3-7 | P01780 | 0.03 | -0.65 |
| Immunoglobulin heavy variable 3-72 | A0A0B4J1Y9 | 0.01 | -0.56 |
| Immunoglobulin heavy variable 3-73 | A0A0B4J1V6 | 0.01 | -0.60 |
| Immunoglobulin heavy variable 4-28 | A0A0C4DH34 | 0.01 | -0.53 |
| Immunoglobulin heavy variable 4-38-2 | P0DP08 | 0.01 | -0.66 |
| Immunoglobulin heavy variable 5-51 | A0A0C4DH38 | 0.03 | -0.45 |
| Immunoglobulin J chain | P01591 | 0.01 | -0.67 |
| Immunoglobulin kappa variable 1-33 | P01594 | 0.01 | -0.32 |
| Immunoglobulin kappa variable 1-5 | P01602 | 0.02 | -0.44 |
| Immunoglobulin kappa variable 2-24 | A0A0C4DH68 | 0.01 | -0.89 |
| Immunoglobulin kappa variable 2-29 | A2NJV5 | 0.02 | -0.71 |
| Immunoglobulin kappa variable 3-20 | P01619 | 0.01 | -0.47 |
| Immunoglobulin kappa variable 4-1 | P06312 | < 0.01 | -0.70 |
| Immunoglobulin lambda variable 1-51 | P01701 | 0.01 | -0.51 |
| Immunoglobulin lambda variable 2-11 | P01706 | 0.01 | -0.78 |
| Immunoglobulin lambda variable 3-19 | P01714 | 0.02 | -0.55 |
| Immunoglobulin lambda variable 4-69 | A0A075B6H9 | 0.02 | -0.74 |
| Immunoglobulin lambda variable 8-61 | A0A075B6I0 | < 0.01 | -0.82 |
| Lymphatic vessel endothelial hyaluronic acid receptor 1 | Q9Y5Y7 | 0.03 | -0.26 |
| Lysosome-associated membrane glycoprotein 2 | P13473 | < 0.01 | -0.31 |
| Phospholipid transfer protein | P55058 | < 0.01 | -0.33 |
| Platelet factor 4 | P02776 | 0.03 | -0.71 |
| Polymeric immunoglobulin receptor | P01833 | 0.01 | -0.53 |
| Probable non-functional immunoglobulin heavy variable 3-35 | A0A0C4DH35 | 0.02 | -0.47 |
| Serotransferrin | P02787 | 0.01 | -0.57 |
| Serum paraoxonase/lactonase 3 | Q15166 | 0.01 | -0.28 |
| Transferrin receptor protein 1 | P02786 | 0.01 | -0.37 |
| ADAMTS-like protein 2 | Q86TH1 | 0.02 | 0.28 |
| Carbonic anhydrase 1 | P00915 | 0.01 | 0.37 |
| Carboxypeptidase Q | Q9Y646 | 0.02 | 0.34 |
| Catalase | P04040 | < 0.01 | 0.35 |
| C-C motif chemokine 14 | Q16627 | 0.05 | 0.38 |
| CD59 glycoprotein | P13987 | 0.01 | 0.26 |
| Ceruloplasmin | P00450 | 0.01 | 0.26 |
| Complement C1r subcomponent-like protein | Q9NZP8 | 0.01 | 0.26 |
| Complement component C6 | P13671 | < 0.0001 | 0.28 |
| Complement component C9 | P02748 | 0.01 | 0.41 |
| Complement factor B | P00751 | < 0.01 | 0.39 |
| Complement factor I | P05156 | 0.01 | 0.25 |
| Inter-alpha-trypsin inhibitor heavy chain H2 | P19823 | < 0.01 | 0.32 |
| Lysosome-associated membrane glycoprotein 1 | P11279 | 0.04 | 0.28 |
| Peroxiredoxin-2 | P32119 | 0.01 | 0.32 |
| Trans-Golgi network integral membrane protein 2 | O43493 | 0.03 | 0.31 |
| Triosephosphate isomerase | P60174 | 0.01 | 0.66 |
| Vinculin | P18206 | 0.02 | 0.30 |
| ZNF511-PRAP1 readthrough | H7BY64 | 0.01 | 0.31 |

Proteins listed with significant differential abundances by pairwise comparisons in limma ("BC\_t0 vs Ctrl"). Proteins are listed by their names with their corresponding UniProt ID, adjusted p-value and fold change, which is expected to be underestimated by 2-fold Ratio-Compression due to TMT-Labeling.

64 % (#27) of these downregulated proteins were immunoglobulin components (Table VII). Looking at the subgroups CR and NR pre-NACT compared to Ctrl, the downregulation of immunoglobulin components compared to the control group is only present in the NR group (Figure 13, Tables VIII and IX).

**Figure 13.**
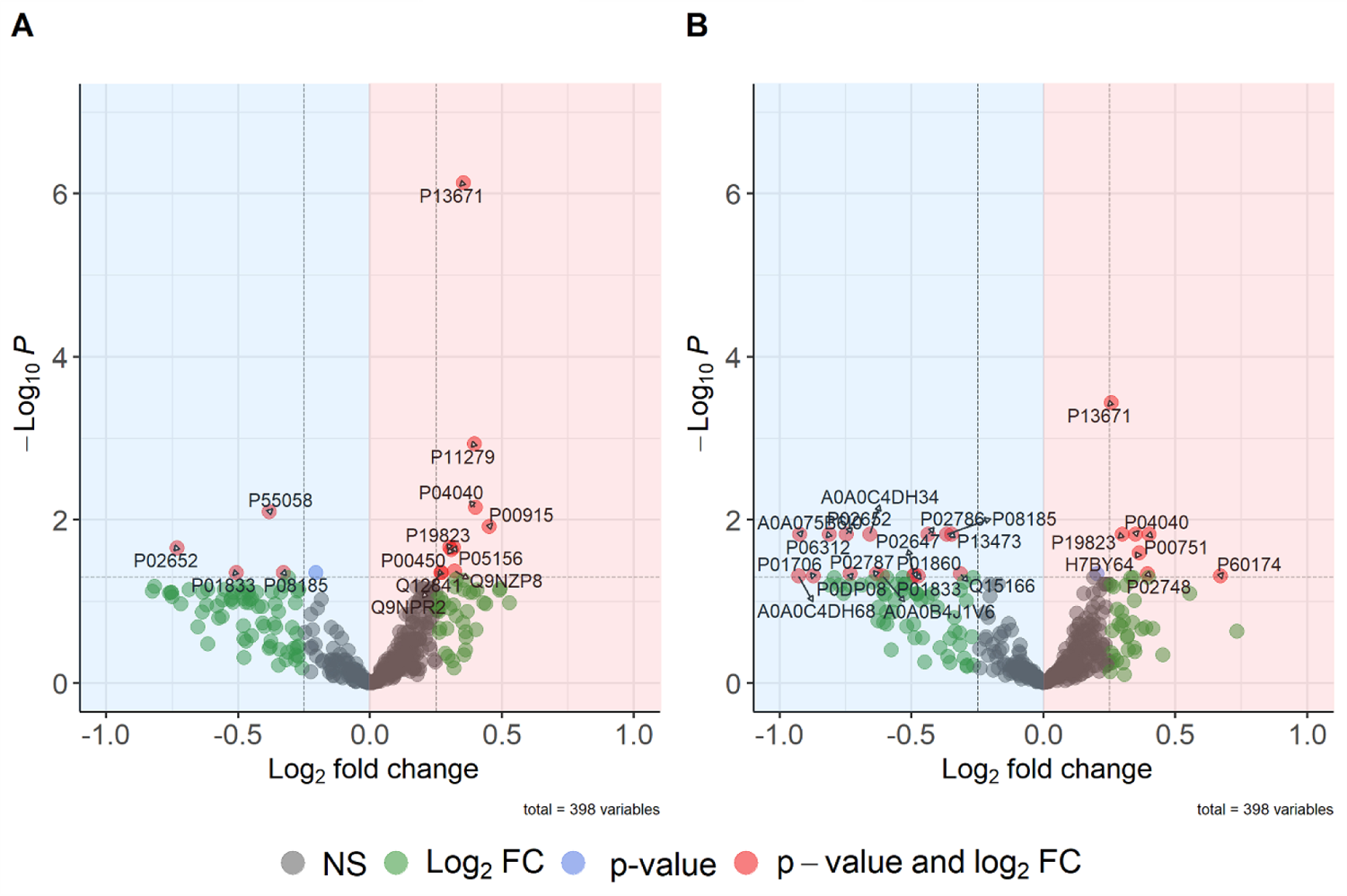
Volcano plots showing the differential protein abundance by pairwise comparisons in limma (“CR_t0 vs. Ctrl, NR_t0 vs. Ctrl”). Protein intensities pre-NACT compared to the control group are shown for CR (A) and NR (B). The log2 fold changes (log2FC) are plotted on the x-axis and the corresponding adjusted p-values in -log10 scale are plotted on the y-axis. Significance was defined by an adjusted p-value < 0.05, log2FC ≥ 0.25 (proteins meeting these criteria are coloured red and marked with their corresponding UniProt ID). Fold change is expected to be underestimated by 2-fold ratio compression due to TMT labelling.

**Table VIII.** Significantly differential regulated proteins pre-NACT in the group of CR compared to the Control Group.

| Protein name | UniProt ID | Adj. p-value | Log2(CR_t0/Ctrl) |
| --- | --- | --- | --- |
| Apolipoprotein A-II | P02652 | 0.02 | -0.73 |
| Corticosteroid-binding globulin | P08185 | 0.04 | -0.33 |
| Phospholipid transfer protein | P55058 | 0.01 | -0.38 |
| Polymeric immunoglobulin receptor | P01833 | 0.04 | -0.51 |
| Carbonic anhydrase 1 | P00915 | 0.01 | 0.45 |
| Catalase | P04040 | 0.01 | 0.40 |
| Ceruloplasmin | P00450 | 0.02 | 0.40 |
| Complement C1r subcomponent-like protein | Q9NZP8 | 0.04 | 0.30 |
| Complement component C6 | P13671 | < 0.0001 | 0.35 |
| Complement factor I | P05156 | 0.02 | 0.32 |
| Follistatin-related protein 1 | Q12841 | 0.04 | 0.27 |
| Inter-alpha-trypsin inhibitor heavy chain H2 | P19823 | 0,02 | 0.30 |
| Lysosome-associated membrane lycoprotein 1 | P11279 | < 0.01 | 0.39 |
| Semaphorin-4B | Q9NPR2 | 0.04 | 0.27 |

Proteins listed with significant differential abundances by pairwise comparisons in limma ("CR\_t0 vs Ctrl"). Proteins are listed by their names with their corresponding UniProt ID, adjusted p-value and fold change, which is expected to be underestimated by 2-fold Ratio-Compression due to TMT-Labeling.

**Table IX.** Significantly differential regulated proteins for NR pre-NACT compared to the Control Group.

| Protein name | UniProt ID | Adj. p-value | Log2(NR_t0/Ctrl) |
| --- | --- | --- | --- |
| Apolipoprotein A-I | P02647 | 0.05 | -0.49 |
| Apolipoprotein A-II | P02652 | 0.01 | -0.75 |
| Corticosteroid-binding globulin | P08185 | 0.01 | -0.37 |
| Immunoglobulin heavy constant gamma 3 | P01860 | 0.05 | -0.49 |
| Immunoglobulin heavy variable 3-73 | A0A0B4J1V6 | 0.05 | -0.61 |
| Immunoglobulin heavy variable 4-28 | A0A0C4DH34 | 0.02 | -0.66 |
| Immunoglobulin heavy variable 4-38-2 | P0DP08 | 0.05 | -0.73 |
| Immunoglobulin kappa variable 2-24 | A0A0C4DH68 | 0.05 | -0.93 |
| Immunoglobulin kappa variable 4-1 | P06312 | 0.01 | -0.81 |
| Immunoglobulin lambda variable 2-11 | P01706 | 0.05 | -0.87 |
| Immunoglobulin lambda variable 8-61 | A0A075B6I0 | 0.01 | -0.92 |
| Lysosome-associated membrane glycoprotein 2 | P13473 | 0.01 | -0.35 |
| Polymeric immunoglobulin receptor | P01833 | 0.05 | -0.47 |
| Serotransferrin | P02787 | 0.05 | -0.63 |
| Serum paraoxonase/lactonase 3 | Q15166 | 0.05 | -0.31 |
| Transferrin receptor protein 1 | P02786 | 0.015 | -0.44 |
| Complement component C6 | P13671 | < 0.001 | 0.26 |
| Catalase | P04040 | 0.01 | 0.35 |
| Complement component C9 | P02748 | 0.05 | 0.39 |
| Complement factor B | P00751 | 0.01 | 0.40 |
| Inter-alpha-trypsin inhibitor heavy chain H2 | P19823 | 0.01 | 0.30 |
| Triosephosphate isomerase | P60174 | 0.05 | 0.67 |
| ZNF511-PRAP1 readthrough | H7BY64 | 0.02 | 0.36 |

Proteins listed with significant differential abundances by pairwise comparisons in limma ("NR\_t0 vs Ctrl"). Proteins are listed by their names with their corresponding UniProt ID, adjusted p-value and fold change, which is expected to be underestimated by 2-fold Ratio-Compression due to TMT-Labeling.

However, the immunoaffinity depletion of Seppro® IgY14 also affects immunoglobulins such as IgG, IgM, IgA. Therefore, this difference in the serum proteome will not be discussed further.

Furthermore, we report a significant downregulation of apolipoprotein AI (P02647, adjusted p-value = 0.01) and AII (P02652, adjusted p-value < 0.01) in BC pre-NACT samples.

We report a significant enrichment of complement components 1RL (Q9NZP8, adjusted p-value = 0.01), 6 (P13671, adjusted p-value = < 0.0001) and 9 (P02748, adjusted p-value = 0.01) in BC pre-NACT samples compared to Ctrl (Figure 12, Table VII). In addition, CD59 (P13987, adjusted p-value = 0.01), a regulator of complement activation known as the MAC inhibitor protein, showed increased intensity in pre-NACT samples.

Inter-alpha trypsin inhibitor heavy chain 2 (ITIH2, P19823, adjusted p-value = 0.01) was upregulated in BC pre-NACT samples compared to Ctrl. ITIH4, one of three or four chains of the inter-alpha trypsin inhibitor proteins, has been described in several studies as a potential marker for the early detection of BC (38). Our results, in conjunction with further studies, may be useful in defining a serum proteomic signature for breast cancer, although this is beyond the scope of this study.

## 4. Conclusion and limitation

In this pilot study, we have successfully profiled the serum proteome response to NACT in breast cancer. Our results indicate that the serum proteome response differs between pCR and non-PCR patients. Further exploratory and confirmatory studies are currently in progress, which may eventually pave the way for liquid biopsy monitoring of NACT. We foresee longitudinal monitoring with individualised baseline values of putative future markers due to the pronounced inter-individual heterogeneity of serum proteome composition (39). Limitations of our study include the size of the cohort, the monocentric design, and the lack of Her2/neu positive patients.

## Acknowledgments

OS acknowledges funding by the Deutsche Forschungsgemeinschaft (DFG, projects 446058856, 466359513, 444936968, 405351425, 431336276, 43198400 (SFB 1453 “NephGen”), 441891347 (SFB 1479 “OncoEscape”), 423813989 (GRK 2606 “ProtPath”), 322977937 (GRK 2344 “MeInBio”)), the ERA PerMed program (BMBF, 01KU1916, 01KU1915A), the German Consortium for Translational Cancer Research (project Impro-Rec), the MatrixCode research group, FRIAS, Freiburg, the investBW program BW1_1198/03, the ERA TransCan program (project 01KT2201,“PREDICO”), and the BMBF KMUi program (project 13GW0603E, project ESTHER). TE acknowledges funding by the Bundesministerium für Bildung und Forschung (project VIP plus “Mammacheck” 03VP08290), Deutsche Krebsgesellschaft (IIT 110536), Fördergesellschaft Forschung Tumorbiologie (1040139401; 1020050201), Gilead Research Grant Breast Cancer. This study was funded by the Fördergesellschaft Forschung Tumorbiologie (funding number:1020050201), Freiburg im Breisgau.

## CRediT authorship contribution statement

**Ines Derya Steenbuck:** Writing - Original Draft preparation, Formal analysis, Visualization, Investigation, Methodology **Miguel Cosenza-Contreras:** Software **Klemens Fröhlich:** Software, Investigation **Bettina Mayer:** Investigation **Tilman Werner:** Methodology **Matthias Fahrner** Methodology **Frank Hause:** Visualization, Software **Adrianna Seredynska:** Investigation **Tobias Feilen:** Investigation **Andrea Ritter:** Data Curation **Armelle Guénégou-Arnoux:** Software **Martin L. Biniossek:** Investigation **Daniela Weiss:** Data Curation **Claudia Nöthling:** Data Curation **Markus Jäger:** Data Curation **Thalia Erbes:** Project administration, Funding acquisition **Oliver Schilling:** Supervision, Writing - Review & Editing, Project administration, Funding acquisition

## Abbreviations

AC: Anthycycline and Cyclophosphamide

BC: Breast Cancer

BCA: Bicinchoninic Acid

CR: Complete Remission

Ctrl: Control Group

LC-MS/MS: Liquid Chromatography-Tandem Mass Spectrometry

NACT: Neoadjuvant Chemotherapy

NR: Non-Complete Remission

P: Paclitaxel

pCR: Pathological Complete Remission

TMT: Tandem Mass Tag

