## Supplemental Figures_Table for "Serum Proteome Profiling Identifies N-Cadherin and C-Met as Early Marker Candidates of Therapeutic Response to Neoadjuvant Chemotherapy in Breast Cancer"

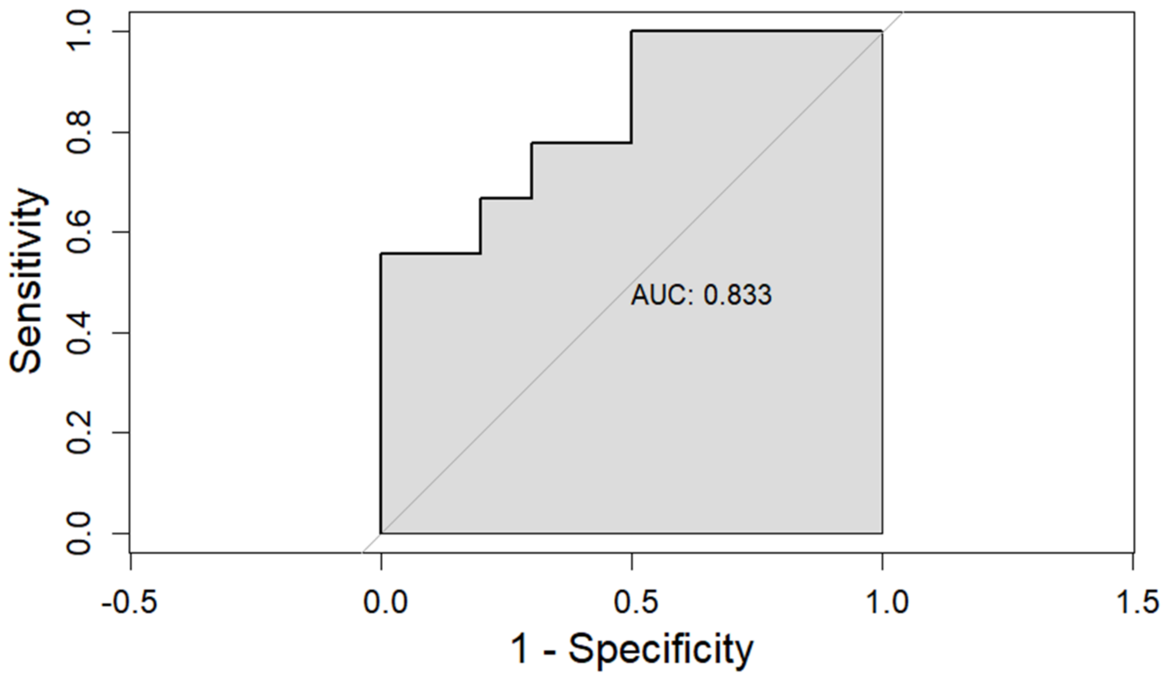


**Figure 1 Supplement** Receiver operating characteristic (ROC) curve for time dependent delta of centrosomal protein, contactin-1, cholinesterase, SHBG

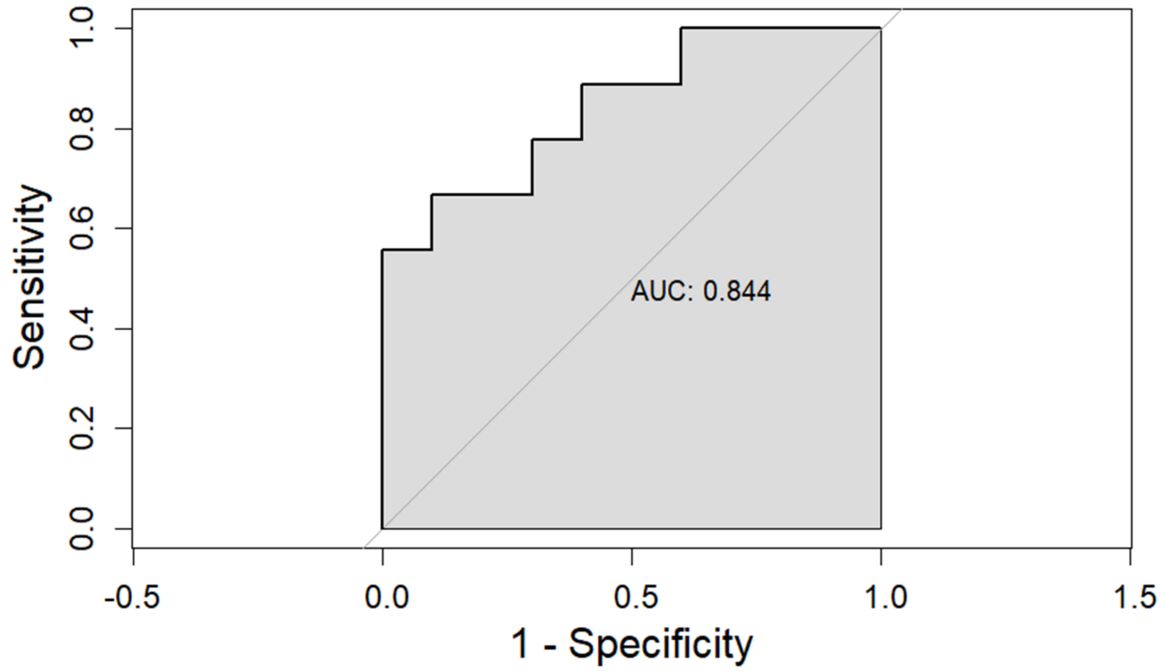


**Figure 2 Supplement** Receiver operating characteristic (ROC) curve for time dependent delta of centrosomal protein, cholinesterase

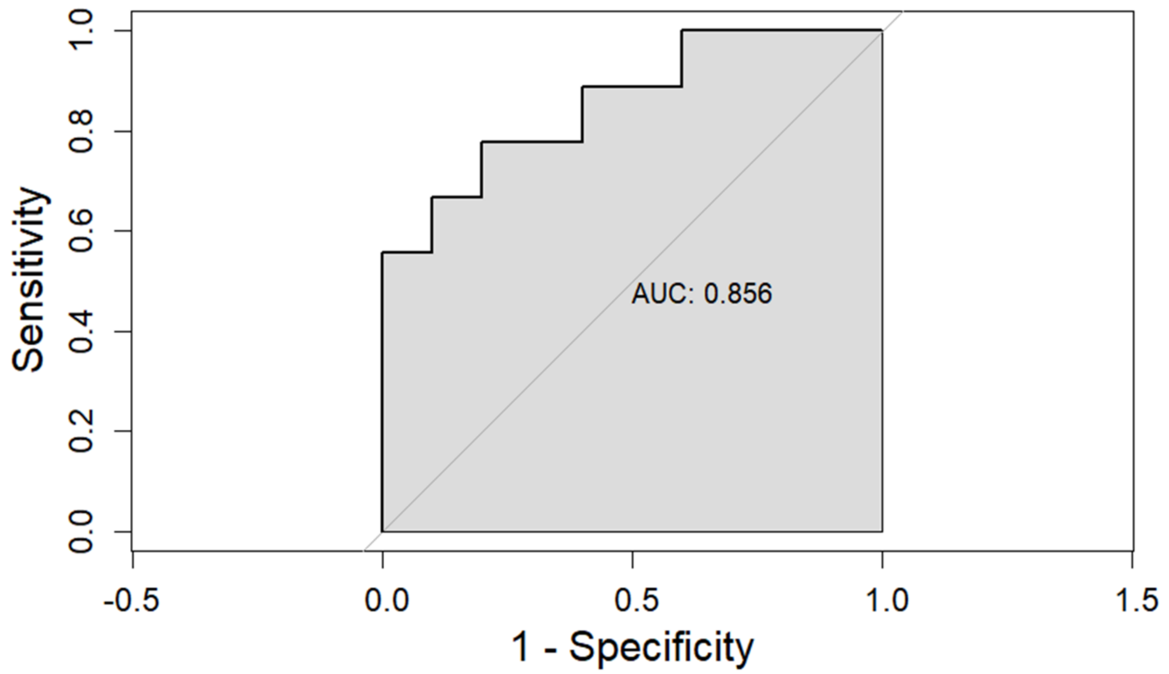


**Figure 3 Supplement** Receiver operating characteristic (ROC) curve for time dependent delta of centrosomal protein

**Supplement Table 1**

Identified proteins with no missing values over all samples

| UniProt ID | Protein name |
|---|---|
| P08195 | 4F2 cell-surface antigen heavy chain |
| Q76LX8 | A disintegrin and metalloproteinase with thrombospondin motifs 13 |
| P63261 | Actin, cytoplasmic 2 |
| Q86TH1 | ADAMTS-like protein 2 |
| Q6UY14 | ADAMTS-like protein 4 |
| P01011 | Alpha-1-antichymotrypsin |
| P01009 | Alpha-1-antitrypsin |
| P04217 | Alpha-1B-glycoprotein |
| P06733 | Alpha-enolase |
| Q16706 | Alpha-mannosidase 2 |
| P03950 | Angiogenin |

| | |
|---|---|
| P12821 | Angiotensin-converting enzyme |
| P01019 | Angiotensinogen |
| O43866 | antigen-like OS=Homo sapiens |
| P02647 | Apolipoprotein A-I |
| P02652 | Apolipoprotein A-II |
| P02654 | Apolipoprotein C-I |
| P02656 | Apolipoprotein C-III |
| P55056 | Apolipoprotein C-IV |
| Q13790 | Apolipoprotein F |
| O14791 | Apolipoprotein L1 |
| O95445 | Apolipoprotein M |
| P08519 | Apolipoprotein(a) |
| P61769 | Beta-2-microglobulin |
| Q96KN2 | Beta-Ala-His dipeptidase |
| P04003 | C4b-binding protein alpha chain |
| P12830 | Cadherin-1 |
| P55290 | Cadherin-13 |
| P19022 | Cadherin-2 |
| P33151 | Cadherin-5 |
| Q9HBB8 | Cadherin-related family member 5 |
| P00915 | Carbonic anhydrase 1 |
| P00918 | Carbonic anhydrase 2 |
| Q96IY4 | Carboxypeptidase B2 |
| P15169 | Carboxypeptidase N catalytic chain |

| | |
|---|---|
| Q9Y646 | Carboxypeptidase Q |
| Q9NQ79 | Cartilage acidic protein 1 |
| P49747 | Cartilage oligomeric matrix protein |
| P25774 | Cathepsin S |
| P11717 | Cation-independent mannose-6-phosphate receptor |
| Q16627 | C-C motif chemokine 14 |
| P55774 | C-C motif chemokine 18 |
| Q6YHK3 | CD109 antigen |
| Q13740 | CD166 antigen |
| P16070 | CD44 antigen |
| P13987 | CD59 glycoprotein |
| Q9BY67 | Cell adhesion molecule 1 |
| Q8TEP8 | Centrosomal protein of 192 kDa |
| Q9NTU7 | Cerebellin-4 |
| P06276 | Cholinesterase OS=Homo sapiens |
| P00740 | Coagulation factor IX |
| P00748 | Coagulation factor XII |
| P00488 | Coagulation factor XIII A chain |
| P23528 | Cofilin-1 |
| P12109 | Collagen alpha-1(VI) chain |
| P39060 | Collagen alpha-1(XVIII) chain |
| P12111 | Collagen alpha-3(VI) chain |
| P02747 | Complement C1q subcomponent subunit C |
| P00736 | Complement C1r subcomponent |

| | |
|---|---|
| P09871 | Complement C1s subcomponent |
| P06681 | Complement C2 |
| P01024 | Complement C3 |
| P01031 | Complement C5 |
| P13671 | Complement component C6 |
| P10643 | Complement component C7 |
| P07357 | Complement component C8 alpha chain |
| P07358 | Complement component C8 beta chain |
| P07360 | Complement component C8 gamma chain |
| P02748 | Complement component C9 |
| P00751 | Complement factor B |
| P08603 | Complement factor H |
| Q03591 | Complement factor H-related protein 1 |
| P36980 | Complement factor H-related protein 2 |
| Q02985 | Complement factor H-related protein 3 |
| Q92496 | Complement factor H-related protein 4 |
| Q9BXR6 | Complement factor H-related protein 5 |
| P05156 | Complement factor I |
| P20023 | Complement receptor type 2 |
| Q8IWV2 | Contactin-4 |
| P08185 | Corticosteroid-binding globulin |
| P01034 | Cystatin-C OS=Homo sapiens |
| P15924 | Desmoplakin |
| Q01459 | Di-N-acetylchitobiase |

| | |
|---|---|
| Q14118 | Dystroglycan |
| Q12805 | EGF-containing fibulin-like extracellular matrix protein 1 |
| P17813 | Endoglin |
| Q16610 | Extracellular matrix protein 1 |
| P08294 | Extracellular superoxide dismutase [Cu-Zn] |
| Q9UGM5 | Fetuin-B |
| P35555 | Fibrillin-1 |
| P02675 | Fibrinogen beta chain |
| P02679 | Fibrinogen gamma chain |
| P11362 | Fibroblast growth factor receptor 1 |
| Q86WI1 | Fibrocystin-L |
| P02751 | Fibronectin |
| P23142 | Fibulin-1 |
| O75636 | Ficolin-3 OS=Homo sapiens |
| P21333 | Filamin-A |
| Q12841 | Follistatin-related protein 1 |
| P16930 | Fumarylacetoacetase |
| P17931 | Galectin-3 |
| Q92820 | Gamma-glutamyl hydrolase |
| P06396 | Gelsolin |
| P22352 | Glutathione peroxidase 3 |
| P00390 | Glutathione reductase, mitochondrial |
| P48637 | Glutathione synthetase |
| P04406 | Glyceraldehyde-3-phosphate dehydrogenase |

| | |
|---|---|
| P00739 | Haptoglobin-related protein |
| P68871 | Hemoglobin subunit beta |
| P02790 | Hemopexin |
| P08581 | Hepatocyte growth factor receptor |
| P26927 | Hepatocyte growth factor-like protein |
| Q14520 | Hyaluronan-binding protein 2 |
| Q9Y6R7 | IgGFc-binding protein |
| P01859 | Immunoglobulin heavy constant gamma 2 |
| P01860 | Immunoglobulin heavy constant gamma 3 |
| P01871 | Immunoglobulin heavy constant mu |
| P23083 | Immunoglobulin heavy variable 1-2 |
| A0A0B4J1V2 | Immunoglobulin heavy variable 2-26 |
| P01817 | Immunoglobulin heavy variable 2-5 |
| A0A0C4DH43 | Immunoglobulin heavy variable 2-70D |
| A0A075B7B8 | Immunoglobulin heavy variable 3/OR16-12 (non-functional) |
| A0A0B4J1V0 | Immunoglobulin heavy variable 3-15 |
| A0A0C4DH35 | Immunoglobulin heavy variable 3-35 |
| P01780 | Immunoglobulin heavy variable 3-7 |
| A0A0B4J1V6 | Immunoglobulin heavy variable 3-73 |
| A0A0C4DH34 | Immunoglobulin heavy variable 4-28 |
| P0DP08 | Immunoglobulin heavy variable 4-38-2 |
| A0A0C4DH38 | Immunoglobulin heavy variable 5-51 |
| P01834 | Immunoglobulin kappa constant |
| P01594 | Immunoglobulin kappa variable 1-33 |

| | |
|---|---|
| P01602 | Immunoglobulin kappa variable 1-5 |
| A0A0C4DH68 | Immunoglobulin kappa variable 2-24 |
| A2NJV5 | Immunoglobulin kappa variable 2-29 |
| P06310 | Immunoglobulin kappa variable 2-30 |
| P04433 | Immunoglobulin kappa variable 3-11 |
| P01619 | Immunoglobulin kappa variable 3-20 |
| P06312 | Immunoglobulin kappa variable 4-1 |
| A0M8Q6 | Immunoglobulin lambda constant 7 |
| P01700 | Immunoglobulin lambda variable 1-47 |
| P01701 | Immunoglobulin lambda variable 1-51 |
| P01714 | Immunoglobulin lambda variable 3-19 |
| A0A075B6H9 | Immunoglobulin lambda variable 4-69 |
| A0A075B6I0 | Immunoglobulin lambda variable 8-61 |
| P01344 | Insulin-like growth factor II |
| P18065 | Insulin-like growth factor-binding protein 2 |
| P17936 | Insulin-like growth factor-binding protein 3 |
| P24593 | Insulin-like growth factor-binding protein 5 |
| P24592 | Insulin-like growth factor-binding protein 6 |
| Q16270 | Insulin-like growth factor-binding protein 7 |
| P05556 | Integrin beta-1 |
| P19827 | Inter-alpha-trypsin inhibitor heavy chain H1 |
| Q06033 | Inter-alpha-trypsin inhibitor heavy chain H3 |
| Q14624 | Inter-alpha-trypsin inhibitor heavy chain H4 |
| P13598 | Intercellular adhesion molecule 2 |

| | |
|---|---|
| P32942 | Intercellular adhesion molecule 3 |
| P29622 | Kallistatin |
| Q5T749 | Keratinocyte proline-rich protein |
| P02788 | Lactotransferrin |
| Q14766 | Latent-transforming growth factor beta-binding protein 1 |
| P02750 | Leucine-rich alpha-2-glycoprotein |
| Q13449 | Limbic system-associated membrane protein |
| P18428 | Lipopolysaccharide-binding protein |
| P00338 | L-lactate dehydrogenase A chain |
| P07195 | L-lactate dehydrogenase B chain |
| P14151 | L-selectin |
| P51884 | Lumican |
| Q9Y5Y7 | Lymphatic vessel endothelial hyaluronic acid receptor 1 |
| P11279 | Lysosome-associated membrane glycoprotein 1 |
| P13473 | Lysosome-associated membrane glycoprotein 2 |
| P61626 | Lysozyme C |
| P22897 | Macrophage mannose receptor 1 |
| P48740 | Mannan-binding lectin serine protease 1 |
| P11226 | Mannose-binding protein C |
| P33908 | Mannosyl-oligosaccharide 1,2-alpha-mannosidase IA |
| P10721 | Mast/stem cell growth factor receptor Kit |
| Q16853 | Membrane primary amine oxidase |
| P01033 | Metalloproteinase inhibitor 1 |
| P16035 | Metalloproteinase inhibitor 2 |

| | |
|---|---|
| P08571 | Monocyte differentiation antigen CD14 |
| Q13201 | Multimerin-1 |
| Q9H8L6 | Multimerin-2 |
| Q7Z7M0 | Multiple epidermal growth factor-like domains protein 8 |
| P05164 | Myeloperoxidase |
| Q99972 | Myocilin |
| Q92859 | Neogenin |
| P13591 | Neural cell adhesion molecule 1 |
| O15394 | Neural cell adhesion molecule 2 |
| O00533 | Neural cell adhesion molecule L1-like protein |
| Q92823 | Neuronal cell adhesion molecule |
| P80188 | Neutrophil gelatinase-associated lipocalin |
| P14543 | Nidogen-1 |
| Q99784 | Noelin |
| P61916 | NPC intracellular cholesterol transporter 2 |
| P32119 | Peroxiredoxin-2 |
| Q96S96 | Phosphatidylethanolamine-binding protein 4 |
| P80108 | Phosphatidylinositol-glycan-specific phospholipase D |
| P55058 | Phospholipid transfer protein |
| P36955 | Pigment epithelium-derived factor |
| P05155 | Plasma protease C1 inhibitor |
| P05154 | Plasma serine protease inhibitor |
| P13796 | Plastin-2 |
| P02775 | Platelet basic protein |

| | |
|---|---|
| Q5VY43 | Platelet endothelial aggregation receptor 1 |
| P02776 | Platelet factor 4 |
| P10720 | Platelet factor 4 variant |
| P40197 | Platelet glycoprotein V |
| O00592 | Podocalyxin |
| P15151 | Poliovirus receptor |
| P20742 | Pregnancy zone protein |
| Q9UHG3 | Prenylcysteine oxidase 1 |
| Q15113 | Procollagen C-endopeptidase enhancer 1 |
| P27918 | Properdin |
| P06702 | Protein S100-A9 |
| Q92954 | Proteoglycan 4 |
| P10586 | Receptor-type tyrosine-protein phosphatase F |
| P34096 | Ribonuclease 4 |
| P07998 | Ribonuclease pancreatic |
| Q86VB7 | Scavenger receptor cysteine-rich type 1 protein M130 |
| Q13103 | Secreted phosphoprotein 24 |
| P49908 | Selenoprotein P |
| Q9NPR2 | Semaphorin-4B |
| P10124 | Serglycin |
| P02787 | Serotransferrin |
| P35542 | Serum amyloid A-4 protein |
| P02743 | Serum amyloid P-component |
| P27169 | Serum paraoxonase/arylesterase 1 |

| | |
|---|---|
| Q15166 | Serum paraoxonase/lactonase 3 |
| P04278 | Sex hormone-binding globulin |
| A1L4H1 | Soluble scavenger receptor cysteine-rich domain-containing protein SSC5D |
| P09486 | SPARC |
| Q14515 | SPARC-like protein 1 |
| P00441 | Superoxide dismutase [Cu-Zn] |
| Q6UWP8 | Suprabasin |
| Q7Z7G0 | Target of Nesh-SH3 |
| P24821 | Tenascin |
| P22105 | Tenascin-X |
| P10599 | Thioredoxin |
| P07996 | Thrombospondin-1 |
| P05543 | Thyroxine-binding globulin |
| P04066 | Tissue alpha-L-fucosidase |
| Q15582 | Transforming growth factor-beta-induced protein ig-h3 |
| O43493 | Trans-Golgi network integral membrane protein |
| P60174 | Triosephosphate isomerase |
| O75382 | Tripartite motif-containing protein 3 |
| P19320 | Vascular cell adhesion protein 1 |
| P35916 | Vascular endothelial growth factor receptor 3 |
| Q6EMK4 | Vasorin |
| Q00341 | Vigilin |
| P18206 | Vinculin |
| P04070 | Vitamin K-dependent protein C |

| | |
|---|---|
| P07225 | Vitamin K-dependent protein S |
| P22891 | Vitamin K-dependent protein Z |
| P54289 | Voltage-dependent calcium channel subunit alpha-2/delta-1 |
| P12955 | Xaa-Pro dipeptidase |
| H7BY64 | ZNF511-PRAP1 readthrough |
| Q15942 | Zyxin |
| P08253 | 72 kDa type IV collagenase |
| Q15848 | Adiponectin |
| Q10588 | ADP-ribosyl cyclase/cyclic ADP-ribose hydrolase 2 |
| P43652 | Afamin |
| P02763 | Alpha-1-acid glycoprotein 1 |
| P19652 | Alpha-1-acid glycoprotein 2 |
| P08697 | Alpha-2-antiplasmin |
| P02765 | Alpha-2-HS-glycoprotein |
| P01023 | Alpha-2-macroglobulin |
| P15144 | Aminopeptidase N |
| Q9Y5C1 | Angiopoietin-related protein 3 |
| P03973 | Antileukoproteinase |
| P01008 | Antithrombin-III |
| P06727 | Apolipoprotein A-IV |
| P04114 | Apolipoprotein B-100 |
| P02655 | Apolipoprotein C-II |
| P05090 | Apolipoprotein D |
| P02649 | Apolipoprotein E |

| | |
|---|---|
| O75882 | Attractin |
| P98160 | Basement membrane-specific heparan sulfate proteoglycan core protein |
| P02749 | Beta-2-glycoprotein 1 |
| P43251 | Biotinidase |
| P80723 | Brain acid soluble protein 1 |
| P20851 | C4b-binding protein beta chain |
| P22792 | Carboxypeptidase N subunit 2 |
| P04040 | Catalase |
| P07339 | Cathepsin D |
| Q9UBR2 | Cathepsin Z |
| P13501 | C-C motif chemokine 5 |
| P43121 | Cell surface glycoprotein MUC18 |
| P00450 | Ceruloplasmin |
| P10909 | Clusterin |
| P12259 | Coagulation factor V |
| P00742 | Coagulation factor X |
| P03951 | Coagulation factor XI |
| P05160 | Coagulation factor XIII B chain |
| P02452 | Collagen alpha-1(I) chain |
| P02745 | Complement C1q subcomponent subunit A |
| P02746 | Complement C1q subcomponent subunit B |
| Q9NZP8 | Complement C1r subcomponent-like protein |
| P0C0L4 | Complement C4-A |
| P0C0L5 | Complement C4-B |

| | |
|---|---|
| Q9NPY3 | Complement component C1q receptor |
| P00746 | Complement factor D |
| Q12860 | Contactin-1 |
| Q9P232 | Contactin-3 |
| P02741 | C-reactive protein |
| P54108 | Cysteine-rich secretory protein 3 |
| P81605 | Dermcidin |
| Q08554 | Desmocollin-1 |
| P27487 | Dipeptidyl peptidase 4 |
| P09172 | Dopamine beta-hydroxylase |
| Q13822 | Ectonucleotide pyrophosphatase/phosphodiesterase family member 2 |
| P11021 | Endoplasmic reticulum chaperone BiP |
| Q9UNN8 | Endothelial protein C receptor |
| P02671 | Fibrinogen alpha chain |
| P04075 | Fructose-bisphosphate aldolase A |
| P05062 | Fructose-bisphosphate aldolase B |
| Q08380 | Galectin-3-binding protein |
| P00738 | Haptoglobin |
| P05546 | Heparin cofactor 2 |
| Q04756 | Hepatocyte growth factor activator |
| P04196 | Histidine-rich glycoprotein |
| Q9Y4L1 | Hypoxia up-regulated protein 1 |
| P01876 | Immunoglobulin heavy constant alpha 1 |
| P01877 | Immunoglobulin heavy constant alpha 2 |

| | |
|---|---|
| P01880 | Immunoglobulin heavy constant delta |
| P01857 | Immunoglobulin heavy constant gamma 1 |
| P01861 | Immunoglobulin heavy constant gamma 4 |
| A0A075B7D0 | Immunoglobulin heavy variable 1/OR15-1 (non-functional) |
| P01742 | Immunoglobulin heavy variable 1-69 |
| S4R460 | Immunoglobulin heavy variable 3/OR16-9 (non-functional) |
| P01766 | Immunoglobulin heavy variable 3-13 |
| P0DP04 | Immunoglobulin heavy variable 3-43D |
| A0A0J9YX35 | Immunoglobulin heavy variable 3-64D |
| A0A0B4J1Y9 | Immunoglobulin heavy variable 3-72 |
| P01591 | Immunoglobulin J chain |
| A0A087WW87 | Immunoglobulin kappa variable 2-40 |
| A0A0C4DH25 | Immunoglobulin kappa variable 3D-20 |
| P0DOY2 | Immunoglobulin lambda constant 2 |
| P01706 | Immunoglobulin lambda variable 2-11 |
| A0A075B6I9 | Immunoglobulin lambda variable 7-46 |
| P15814 | Immunoglobulin lambda-like polypeptide 1 |
| B9A064 | Immunoglobulin lambda-like polypeptide 5 |
| P05019 | Insulin-like growth factor I |
| P22692 | Insulin-like growth factor-binding protein 4 |
| P35858 | Insulin-like growth factor-binding protein complex acid labile subunit |
| P19823 | Inter-alpha-trypsin inhibitor heavy chain H2 |
| P05362 | Intercellular adhesion molecule 1 |
| Q9NPH3 | Interleukin-1 receptor accessory protein |

| | |
|---|---|
| P01042 | Kininogen-1 |
| P24043 | Laminin subunit alpha-2 |
| P07942 | Laminin subunit beta-1 |
| O14960 | Leukocyte cell-derived chemotaxin-2 |
| P07333 | Macrophage colony-stimulating factor 1 receptor |
| O00187 | Mannan-binding lectin serine protease 2 |
| P14780 | Matrix metalloproteinase-9 |
| P26038 | Moesin |
| Q9H1U4 | Multiple epidermal growth factor-like domains protein 9 |
| Q9UNW1 | Multiple inositol polyphosphate phosphatase 1 |
| P20933 | N(4)-(beta-N-acetylglucosaminyl)-L-asparaginase |
| Q96PD5 | N-acetylmuramoyl-L-alanine amidase |
| Q8NF91 | Nesprin-1 |
| O14786 | Neuropilin-1 |
| O95497 | Pantetheinase |
| Q6UXB8 | Peptidase inhibitor 16 |
| P04180 | Phosphatidylcholine-sterol acyltransferase |
| P03952 | Plasma kallikrein |
| P00747 | Plasminogen |
| P07359 | Platelet glycoprotein Ib alpha chain |
| Q6UX71 | Plexin domain-containing protein 2 |
| P01833 | Polymeric immunoglobulin receptor |
| P07737 | Profilin-1 |
| Q07954 | Prolow-density lipoprotein receptor-related protein 1 |

| | |
|---|---|
| P41222 | Prostaglandin-H2 D-isomerase |
| P02760 | Protein AMBP |
| Q9ULI3 | Protein HEG homolog 1 |
| P05109 | Protein S100-A8 |
| Q9UK55 | Protein Z-dependent protease inhibitor |
| P00734 | Prothrombin |
| Q9HBR0 | Putative sodium-coupled neutral amino acid transporter 10 |
| Q12913 | Receptor-type tyrosine-protein phosphatase eta |
| P23470 | Receptor-type tyrosine-protein phosphatase gamma |
| P02753 | Retinol-binding protein 4 |
| Q8WZ75 | Roundabout homolog 4 |
| Q9NQ38 | Serine protease inhibitor Kazal-type 5 |
| Q86U17 | Serpin A11 |
| Q9H299 | SH3 domain-binding glutamic acid-rich-like protein 3 |
| O00391 | Sulfhydryl oxidase 1 |
| P05452 | Tetranectin |
| P02786 | Transferrin receptor protein 1 |
| P29401 | Transketolase |
| Q14956 | Transmembrane glycoprotein NMB |
| P02766 | Transthyretin |
| Q86YW5 | Trem-like transcript 1 protein |
| A0A087WW49 | Uncharacterized protein |
| A0A0J9YY99 | Uncharacterized protein (Fragment) |
| P02774 | Vitamin D-binding protein |

| | |
|---|---|
| P04004 | Vitronectin |
| P04275 | von Willebrand factor |
| P25311 | Zinc-alpha-2-glycoprotein |
